# Interferon-α subtype treatment induces the repression of SRSF1 in HIV-1 target cells and affects HIV-1 post integration steps

**DOI:** 10.1101/2021.06.11.448031

**Authors:** Helene Sertznig, Fabian Roesmann, Barbara Bleekmann, Carina Elsner, Mario Santiago, Jonas Schuhenn, Yvonne Benatzy, Ryan Snodgrass, Stefan Esser, Kathrin Sutter, Ulf Dittmer, Marek Widera

**Author notes:** Correspondence: Dr. rer. nat. Marek Widera, Institute for Medical Virology, University Hospital Frankfurt, Paul-Ehrlich-Str. 40, 60596 Frankfurt am Main.

## Abstract

Efficient replication of HIV-1 depends on balanced levels of host cell components, including cellular splicing factors. Type I interferons (IFN-I), playing a crucial role in the innate immune defense against viral infections, are well known to induce the transcription of IFN-stimulated genes (ISGs) including potent host restriction factors. Not so well known is, that IFN-repressed genes (IRepGs) also affect viral infections by downregulating host dependency factors that are essential for viral replication. So far, knowledge about IRepGs involved in HIV-1 infection is very limited. Here, we demonstrate that expression levels of the serine/arginine-rich splicing factor 1 (SRSF1) were repressed upon treatment with IFNα subtypes in HIV-1 susceptible cell lines as well as primary cells. Furthermore, we could demonstrate in two independent patient cohorts that HIV-1 infection and the concomitant inflammation during the acute and chronic phase, resulted in the strong induction of ISGs, but at the same time significantly repressed SRSF1. 4sU-labeling of newly transcribed mRNAs revealed that IFN-mediated repression of SRSF1 originated from a transcriptional shutdown. Experimental downregulation as well as overexpression of SRSF1 expression levels resulted in crucial changes in HIV-1 LTR-transcription, alternative splice site usage and virus production. While lower SRSF1 levels resulted in low *vif* mRNA levels and thus severely reduced viral infectivity, higher levels of SRSF1 impaired LTR-Tat-activity and HIV-1 particle production.

Our data highlight the so far undescribed role of SRSF1 acting as an IFN-repressed cellular dependency factor decisively regulating HIV-1 post integration steps.

**Author Summary:** IFN-I play a central role in the innate immune defense against viral infections by regulating the expression of interferon stimulated genes (ISGs) and interferon repressed genes (IRepGs). The stimulation of host restriction factors and the reduction of host dependency factors decisively affects the efficiency of HIV-1 replication. After the stable integration of the provirus into the host chromosome, HIV-1 exploits the host cell transcription and splicing machinery for its replication. A network of conserved splice sites and splicing regulatory elements maintain balanced levels of viral transcripts essential for virus production and immune evasion.

We demonstrate the so far undescribed role of the splicing factor SRSF1 as an IRepG crucially involved in HIV-1 RNA processing. In HIV-1 infected individuals, we observed inversely proportional expression of high ISG15 and low SRSF1 levels, which were restored in ART treated patients. We could demonstrate, that IFN-I stimulation of HIV-1 target cells resulted in a significant repression of SRSF1 RNA and protein levels. Since low SRSF1 expression decisively reduced HIV-1 *vif* mRNA levels, a severe impairment of viral replication was observed in APOBEC3G expressing cells. As overexpression negatively affected HIV-1 LTR transcription and virus production, balanced levels of SRSF1 are indispensable for efficient replication.

## Introduction

The human immunodeficiency virus type 1 (HIV-1) depends on cellular components of the host, which are crucial for efficient replication and thus termed host dependency factors (1). Once integrated into the host genome, HIV-1 uses the cellular transcription apparatus and splicing machinery for viral gene expression. Since important regulatory HIV-1 proteins are expressed from spliced intron-less viral mRNAs, cellular splicing factors and splicing regulatory proteins are indispensable for viral replication. Thus, alternative splicing and exploitation of the full range of the cellular splicing code is required to produce balanced levels of all essential viral mRNAs (2, 3). Type I interferons (IFN-I), which amongst others include 12 different IFNα subtypes and IFNβ, play a crucial role in the innate immune defense against viral infections including HIV-1 (4, 5). All IFNα subtypes have been shown to exert distinct biological activities dependent on their binding affinities, receptor avidity or cell type specificity (6-8). In contrast to the clinically used subtype IFNα2, which shows only limited antiviral activity against HIV-1, IFNα14 has proven to be the most potent subtype against HIV-1 (7, 9, 10). After viral sensing via pattern recognition receptors (PRR) like the Toll-like receptors (TLR) or the cytosolic DNA sensor cyclic GMP-AMP synthase (cGAS), transcription and secretion of IFN-I is induced (11, 12). Binding of IFN-I to the IFNα/β-receptor (IFNAR) induces signaling via the JAK/STAT-pathway and leads to the transcription of hundreds of IFN-stimulated genes (ISG), such as host restriction factors or transcription factors, establishing an antiviral state within the cell (11, 13). Among others, prominent members of ISGs with anti-retroviral activity include ISG15 (IFN-stimulated gene 15), APOBEC3G (apolipoprotein B mRNA editing enzyme, catalytic polypeptide-like 3G), tetherin, Mx-2 (Myxovirus resistance-2), SamHD1 (SAM domain and HD domain-containing protein 1) and IFITM1-3 (Interferon-induced transmembrane protein 1-3) (14, 15).

In addition to the well-described induction of ISGs, it has also been shown that IFNs are able to repress the expression of specific genes, termed IFN-repressed genes (IRepG), which in part are essential for viral replication (16, 17). This downregulation might represent a possible defense mechanism of the cell, limiting essential dependency factors for viral replication (16).

The replication strategy of HIV-1 involves the usage of multiple conserved splice donor and acceptor sequences, which in various combinations enable the generation of more than 50 viral transcript isoforms (3, 18). Balanced levels of all transcript isoforms are crucial for efficient virus replication. The stability of the RNA duplex of the cellular U1 small nuclear (sn) RNA and the respective splice donor site defines the intrinsic strength of a specific 5’-splice site (5’-ss) (19, 20). In addition, the polypyrimidine content in the polypyrimidine tract (PPT) determines the intrinsic strength of a splice acceptor or 3’-ss (21). Furthermore, a complex network of splicing regulatory *cis*-elements, localized on the viral pre-mRNA, can be bound by host cell derived *trans*-acting RNA-binding proteins, which decisively regulate the ratio of HIV-1 transcript isoforms (22, 23). The protein family of serine/arginine-rich splicing factors (SRSF) belongs to the large family of RNA binding proteins (24, 25). Members of this protein family are well known to act as cellular splicing factors (24, 26). Depending on the position of their binding region within an exon or intron, they can enhance or repress the usage of a specific splice site (26-28). Two main structural features are conserved among all SR proteins, the protein-interacting RS-domain, which is rich in arginine and serine (RS) dipeptides, and the RNA recognition motif (RRM) (25). The activity of SR proteins is regulated via phosphorylation of the RS-domain through specific SR protein kinases (SRPK) or other kinases like Akt (29). Furthermore, shuttling of SR proteins between cytoplasm and nucleus is dependent on the phosphorylation state of the RS-domain (29). SR proteins are generally characterized by their ability to interact with both RNA and protein structures simultaneously (25).

As the founding member of the SRSF protein family, serine/arginine-rich splicing factor 1 (SRSF1), formerly known as SRp30a or ASF/SF2 (30), was originally identified to promote spliceosomal assembly and pre-mRNA splicing in HeLa cells (31), as well as to regulate alternative splicing of the SV40 pre-mRNA (32). SRSF1 was shown to bind to the *cis*-regulatory elements ESE (exonic splicing enhancer) M1/M2, ESE-GAR and ESE3 in the HIV-1 genome, facilitating the usage of specific splice sites (33-35). While overexpression of SRSF1 resulted in enhanced Vpr, but reduced Tat1, Gag and Env levels (36-38), knockdown increased levels of all viral RNAs indicating an effect on both alternative splicing and LTR transcription (36). Thus, SRSF1 represents a key regulator and host dependency factor important for efficient HIV-1 RNA processing, enabling the emergence of the protein diversity necessary for efficient viral replication.

In this manuscript, we investigated whether the expression levels of SRSF proteins are influenced by HIV-1 infection or IFN stimulation. We found that IFN-I treatment induces the repression of SRSF1 in HIV-1 host cells affecting viral post integration steps. Our findings suggest, that balanced levels of SRSF1 are crucial for efficient HIV-1 replication, as both higher and lower levels led to severe impairments at the level of LTR transcription, alternative splicing or virus production.

## Results

### SRSF1 is significantly downregulated in HIV-1 infected patients

In a previous RNA-sequencing based study we were able to demonstrate that levels of specific host restriction factors, such as tetherin, Mx-2 or APOBEC3G were upregulated in the gut of HIV-1 infected individuals, confirming chronic inflammation (39). In order to analyze whether also host dependency factors were significantly altered upon HIV-1 infection, we compared gene expression levels of chronically HIV-1 infected patients, either naïve or under antiretroviral (ART) therapy, with those of healthy individuals. Focusing on the expression levels of SRSF mRNAs, we found significantly lower levels of *SRSF1, SRSF3, SRSF7* and *SRSF10* mRNA in chronically HIV-1 infected patients when compared to healthy individuals (**Fig 1a**). Furthermore, *SRSF2, SRSF5* and *SRSF8* transcript levels were also lower in this cohort, albeit the difference was not significant (**Fig 1a**). Surprisingly, we observed that gene expression of SRSF is generally restored in patients under ART treatment and transcript levels of *SRSF1* were even significantly higher when compared to healthy donors (**Fig 1b**). Even under ART-treatment, *SRSF3* and *SRSF10* mRNA expression levels were still lower in chronically HIV-1 infected patients in contrast to healthy individuals (**Fig 1b**). Transcript levels of *SRSF2* and *SRSF7* in ART-treated patients were comparable to the levels observed in the healthy control group. (**Fig 1b**). A marginal but significant difference in SRSF expression levels in healthy individuals and HIV-1 infected ART-treated patients was also observed for *SRSF9*, however the total amount of transcripts was low abundant (**Fig 1b**). In all patient groups, *SRSF12* transcript levels were only slightly above the limit of detection (**Fig 1a-b**).

**Fig 1:**
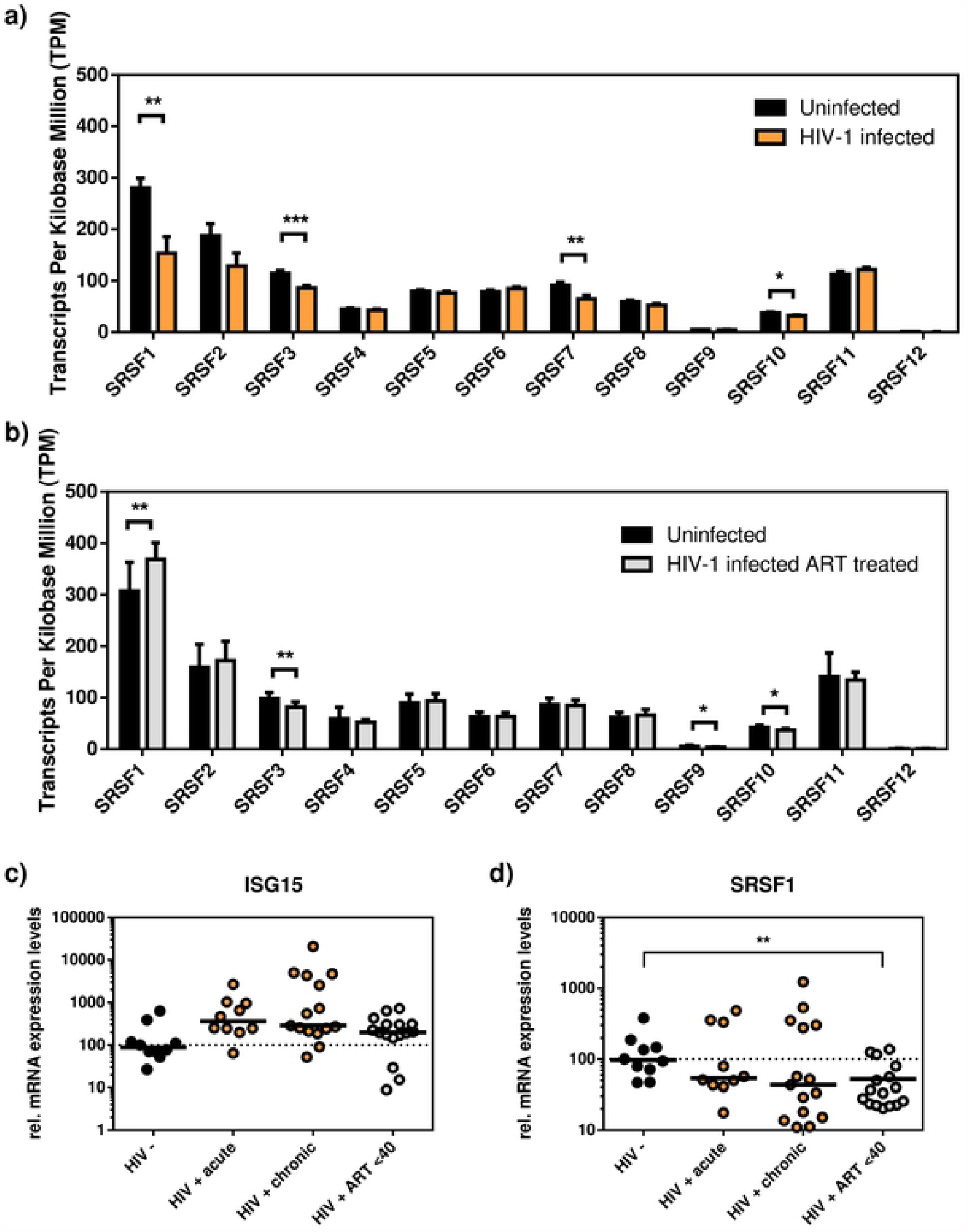
**a) – b) SRSF levels in naïve or ART-treated HIV-1 infected individuals.** Transcript levels of SRSF genes were measured in colonic samples using RNA-sequencing analysis. Comparison of results from **a)** naïve HIV-1 infected and healthy individuals and **b)** ART-treated HIV-1 infected and healthy individuals. Mann-Whitney statistical analysis was performed to determine differences between unmatched groups. **c) – d) SRSF1 levels inversely correlate with ISG15 expression**. RT-qPCR results for the mRNA expression levels of **c)** ISG15 and **d)** SRSF1 in healthy individuals, acutely and chronically HIV-1 infected patients as well as HIV-1 infected ART-treated individuals. ACTB was used as loading control. Unpaired t-tests were calculated to determine whether the difference between the group of samples reached the level of statistical significance (* p<0.05, ** p<0.01 and *** p<0.001).

Since SRSF1 was the most significant of the differentially expressed genes in the patient groups and also described to be crucially involved in HIV-1 RNA processing (36, 40), we analyzed the expression profile of SRSF1 in HIV-1 infected patients at different phases of infection. For this purpose, PBMCs were isolated from HIV-1 positive individuals during acute infection phase (Fiebig I-V), chronic infection phase (Fiebig VI) or chronic infection phase under ART-treatment as well as from HIV-1 negative donors. Total cellular RNA was isolated and subjected to RT-qPCR analysis. To evaluate whether the patient cohort was representative, we analyzed *ISG15* mRNA expression as a surrogate marker for ISG induction. Acutely and chronically HIV-1 infected patients had strongly increased mRNA expression levels of *ISG15* in PBMCs when compared to healthy individuals, indicating a virus-induced IFN signature proportional to the viral load (**S1 Fig)**. HIV-1 infected patients under ART-treatment showed reduced levels of *ISG15* when compared to untreated HIV-1 infected individuals albeit still higher than healthy individuals (**Fig 1c**). The *ISG15* expression data of our representative cohort were in line with previous observations of HIV-1 infection stimulating IFN induction and thus ISGs (41, 42). In order to investigate, whether repression of SRSF1 correlates with ISG induction in HIV-1 infected patients, we performed an SRSF1-specific RT-qPCR. For acutely and chronically HIV-1 infected patients, as well as ART-treated patients, lower levels of *SRSF1* mRNA were detected in contrast to healthy donors (**Fig 1d**). In chronically infected patients, *SRSF1* levels were downregulated in the majority of the patient derived samples. However, in some patients *SRSF1* mRNA levels were increased in contrast to the healthy control group (**Fig 1d**), a finding which can be deduced from the fact that chronically HIV-1 infected patients generally represent a heterogeneous cohort. ART-treated patients represented a more homogeneous group in comparison to acutely or chronically infected patients showing significantly decreased *SRSF1* mRNA levels (**Fig 1d**). In general, high induction of *ISG15* was concomitant with strong repression of *SRSF1* in single individuals. Thus, we discovered a possible interrelation between the downregulation of SRSF1 and the stimulation of ISGs suggesting that SRSF1 might act as an IFN-repressed gene (IRepG).

### The strength of SRF1 repression is IFNα subtype dependent

We previously showed that the 12 different IFNα subtypes exert different anti-viral activities in HIV-1 infection. Their anti-HIV-1 potential correlated with the induction of ISGs known to be anti-viral restriction factors against HIV-1 (7, 9). Thus, we tested all 12 IFNα subtypes (α1, α2, α4, α5, α6, α7, α8, α10, α14, α16, α17, α21) upon their ability to repress *SRSF1* mRNA expression. First, we used a luciferase reporter cell line which harbors the firefly luciferase gene under the control of the IFN-inducible ISRE promotor to evaluate the biological activity of all IFNα subtypes (43). All IFNα subtypes induced an increase in the luminescent signal compared to the PBS-stimulated control except for IFNα1 (**S2 Fig**).

Next, THP-1 monocytic cells were differentiated into macrophage-like cells using phorbol 12-myristate 13-acetate (PMA) and treated with different IFNα subtypes. In general, the intensity of ISG15 induction reflected the intensity of the luminescent signal in the reporter cells, with the exemption of IFNα8 (**Fig 2a and S2 Fig**). Subtypes IFNα2, α4, α6, α10, α14 and α17 induced the strongest *ISG15* mRNA expression with a 50-to 100-fold increase when compared to the unstimulated control. Treatment with IFNα5, α16 and α21 led to a moderate *ISG15* increase of 15-to 30-fold, while treatment with α1, α7 and α8 only led to a slightly enhanced *ISG15* mRNA expression (**Fig 2a**). The strongest repression of *SRSF1* mRNA expression was observed for IFNα10 and α14 with roughly 4-fold and 10-fold respectively **(Fig 2b)**. The subtypes α4, α6, α7, α8, α17 and α21 induced a moderate *SRSF1* mRNA repression of roughly 2-fold, while the subtypes α1, α2, α5 and α16 only induced a weak downregulation of less than 2-fold (**Fig 2b**). The vast majority of the tested IFNα subtypes showed a positive correlation between *ISG15* induction and *SRSF1* repression (**Fig 2c**). However, IFNα14 downregulated *SRSF1* expression disproportionately strong compared to the *ISG15* induction. In contrast, treatment with IFNα2, α4, and α6 led to strong *ISG15* induction but comparatively moderate *SRSF1* repression. In conclusion, for the majority of IFNα subtypes suppression of *SRSF1* mRNA expression positively correlated with ISG induction. Generally, all IFNα subtypes induced a repression of *SRSF1* mRNA, albeit to highly varying extent, with IFNα10 and IFNα14 being the most potent.

**Fig 2:**
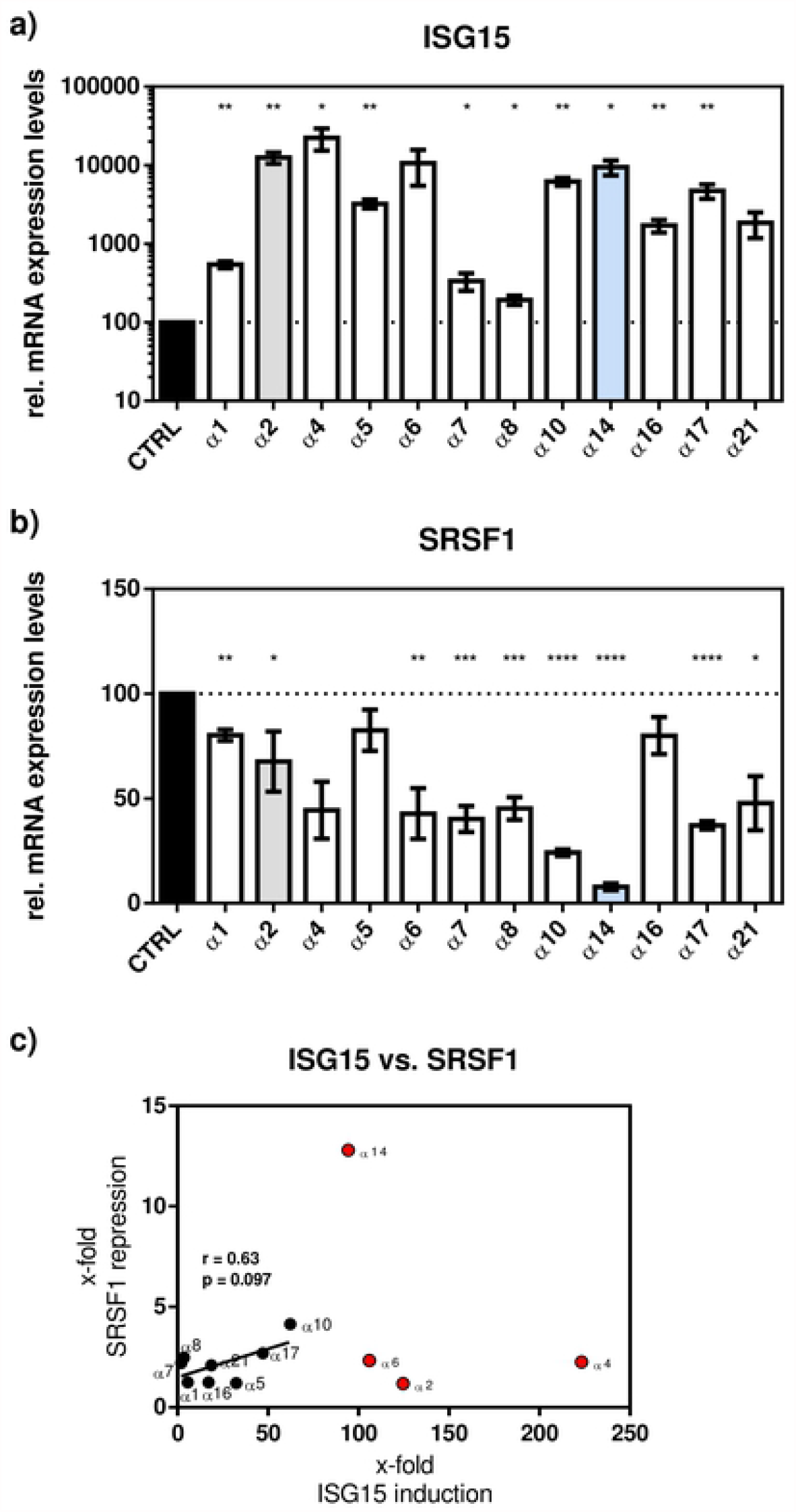
Downregulation of SRSF1 is IFNα subtype dependent. **a) - b)** Differentiated THP-1 cells were treated with the indicated IFN subtype at a concentration of 10 ng/ml. 24 h post treatment, cells were harvested, RNA isolated and subjected to RT-qPCR for measurement of relative **a)** ISG15 and **b)** SRSF1 mRNA expression levels. ACTB was used as loading control. Unpaired t tests were calculated to determine whether the difference between the group of samples reached the level of statistical significance (* p<0.05, ** p<0.01 and *** p<0.001). **c)** Correlation between x-fold repression of SRSF1 mRNA levels and x-fold induction of ISG15 mRNA levels. IFNα subtypes 2, 4, 6 and 14 were excluded from correlation and are marked in red. Pearson correlation coefficient (r) and p-value (p) are indicated.

### SRSF1 is an IRepG in macrophages and T-cells

Next, we examined *SRSF1* expression levels in HIV-1 target cells upon IFN-treatment in a time-course experiment. In our studies, we included IFNα14, the subtype which induced the strongest downregulation of SRSF1 expression levels **(Fig 2)** and is the most potent IFNα subtype against HIV-1 (7). Furthermore, we added IFNα2, which is the IFNα subtype already clinically used for the treatment of other viruses including hepatitis B virus (44). The induction of ISGs was monitored using ISG15.

In differentiated macrophage-like THP-1 cells, we observed a strong 100-to 1000-fold induction of *ISG15* mRNA expression levels after 4 h of treatment with both IFNα2 and IFNα14 (**Fig 3a**). Treatment with IFNα2 resulted in a 13-fold downregulation of *SRSF1* after 12 h while expression levels were restored 24 h post treatment (**Fig 3b**). Treatment with IFNα14 also resulted in a 13-fold downregulation of *SRSF1* after 24 h and a long-lasting effect with a still 6-fold downregulation after 48 h (**Fig 3c**). Overall IFNα14 induced a stronger and more long-lasting repression than IFNα2 **(Fig 3b-c)**.

**Fig 3:**
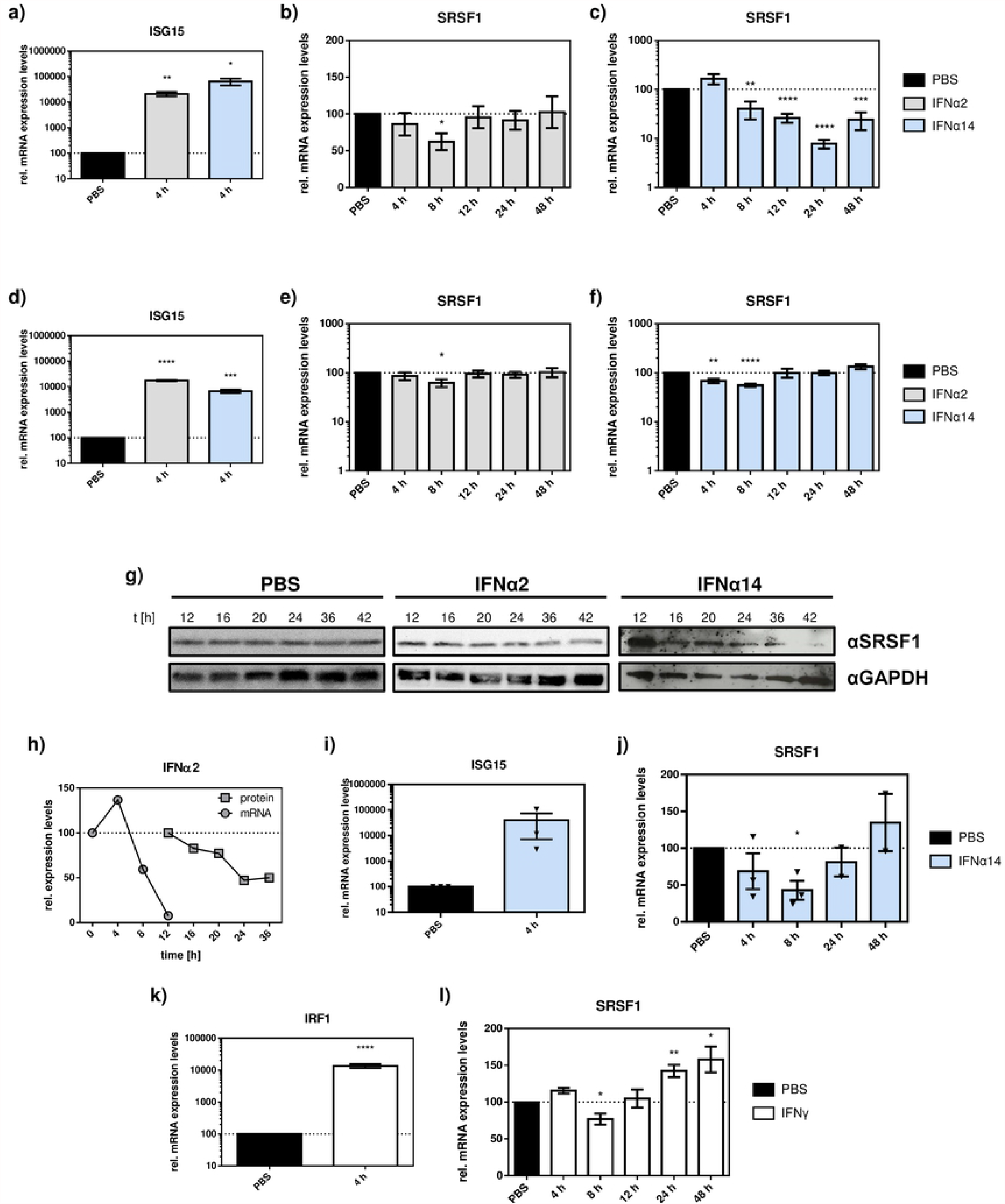
**a) – h) SRSF1 levels in HIV-1 host cells are repressed upon IFN treatment.** Differentiated THP-1 macrophages and Jurkat T-cells were treated with the indicated IFN subtype over a period of 48 h at a concentration of 10 ng/ml before cells were harvested and RNA was isolated. Relative mRNA expression levels of ISG15 and SRSF1 were measured via RT-qPCR. ISG15 mRNA levels were measured after 4 h for **a)** THP-1 cells and **d)** Jurkat cells. SRSF1 mRNA levels were measured at the indicated time points in THP-1 cells after treatment with **b)** IFNα2 and **c)** IFNα14 and in Jurkat cells after treatment with **e)** IFNα2 and **f)** IFNα14. ACTB was used as loading control. Unpaired t-tests were calculated to determine whether the difference between the group of samples reached the level of statistical significance (* p<0.05, ** p<0.01 and *** p<0.001). **g)** Differentiated THP-1 macrophages were treated with IFNα2 or IFNα14 for the indicated amount of time at a concentration of 10 ng/ml before cells were harvested. Proteins were separated by SDS-PAGE, blotted and analyzed with an antibody specific to SRSF1. GAPDH was used as loading control. **h)** Comparison of time-dependent SRSF1-repression after IFNα2-treatment on mRNA and protein level. **i)** – **j) Repression of SRSF1 mRNA levels in primary human macrophages**. Monocyte-derived macrophages (MDMs) were treated with IFNα14 over a period of 48 h at a concentration of 10 ng/ml before cells were harvested and RNA isolated. Relative mRNA expression levels of ISG15 and SRSF1 were measured via RT-qPCR. **i)** ISG15 mRNA levels were measured 4 h post treatment. **j)** SRSF1 mRNA levels were measured at the indicated time points. GAPDH was used as loading control. Unpaired t-tests were calculated to determine whether the difference between the group of samples reached the level of statistical significance (* p<0.05, ** p<0.01 and *** p<0.001). Time points 24 h and 48 h only include two biological replicates. **k) – l) SRSF1 repression in THP-1 cells is type I IFN specific**. Differentiated THP-1 cells were treated with IFNγ over a period of 48 h at a concentration of 10 ng/ml before cells were harvested and RNA was isolated. Relative mRNA expression levels of IRF1 and SRSF1 were measured via RT-qPCR. **k)** IRF1 mRNA levels were measured after 4 h. **l)** SRSF1 mRNA levels were measured at the indicated time points. GAPDH was used as loading control. Unpaired t-tests were calculated to determine whether the difference between the group of samples reached the level of statistical significance (* p<0.05, ** p<0.01 and *** p<0.001).

In Jurkat T-cells, IFN-treatment with both IFNα subtypes led to a strong 10-to 100-fold induction of *ISG15* mRNA expressions levels after 4 h (**Fig 3d**) albeit less pronounced when compared to the THP-1 cells (**Fig 3a**). A time-dependent significant downregulation of *SRSF1* mRNA levels could be observed after treatment with both IFNα2 and IFNα14 (**Fig 3e-f**). Significant downregulation of *SRSF1* in Jurkat T-cells already occurred after 4 to 8 h of treatment and was much less pronounced than in THP-1 cells with an only about 2-fold reduction for both IFNα subtypes (**Fig 3e-f**).

In conclusion, inversely to *ISG15* expression, *SRSF1* was downregulated in HIV-1 target cells, in particular macrophage-like THP-1 cells, upon IFN-I stimulation and thus represents an IRepG. In order to further analyze whether the IFN-induced reduction of SRSF1 can also be observed on the protein level, we performed Western Blot analysis of IFN-treated THP-1 cells. Both treatments with IFNα2 and IFNα14 resulted in a decrease in SRSF1 protein levels 36-42 h post treatment (**Fig 3g**). While treatment with IFNα2 only led to a weak repression, treatment with IFNα14 resulted in a strong downregulation of SRSF1 protein levels (**Fig 3g**), which was in accordance to the results on mRNA expression levels (**Fig 3b-c**). When compared to the mRNA levels, SRSF1 protein levels decrease with a time shift of 12 to 24 h, which might be explained by the half-life of persisting mRNA and protein levels **(Fig 3h)**.

In order to assess whether these findings can be translated to primary human cells, we analyzed gene expression of SRSF1 after treatment with IFNα14 in primary human monocyte-derived macrophages (MDMs). A strong 50-to 500-fold induction was observed for *ISG15* mRNA expression levels after 4 h of treatment **(Fig 3i)**. Concomitantly, a time-dependent repression of SRSF1 was detected with a significantly >2-fold downregulation of *SRSF1* mRNA levels after 8 h **(Fig 3j)**. Although less pronounced than in the cell culture system, IFN-mediated repression of *SRSF1* mRNA expression could thus be confirmed in primary human cells.

To assess whether the downregulation of SRSF1 was IFN-I specific, we included IFNγ as the only member of the type II IFN family (45). Since IFNγ binds to the IFNγ receptor (IFNGR) and activates a distinct signaling pathway (46), the IFN-regulatory factor 1 (IRF1) was chosen as a control of IFN-II specific activation of the gamma interferon activation site (GAS) regulated promotor (47). We used THP-1 macrophage-like cells since they showed the strongest repression of SRSF1 upon IFN-I treatment (**Fig 3b, c, e and f**).

Treatment with IFNγ led to a strong 100-fold induction of *IRF1* after 4 h (**Fig 3k**), but only a weak (1.3-fold) reduction in *SRSF1* mRNA expression levels was detected after 8 h of IFNγ-treatment (**Fig 3l**). However, the overall changes in mRNA expression were negligible when compared to the effect of IFN-I treatment (**Fig 3b-c**). Additionally, a time-dependent increase in *SRSF1* mRNA expression was observed after IFNγ-treatment between 12 h to 48 h, resulting in significantly elevated levels at 24 h and 48 h (**Fig 3l**). Overall, the repression of SRSF1 seems to be a more IFN-I specific effect.

### SRSF1 downregulation occurs on transcriptional level

To further investigate, whether the downregulation of SRSF1 occurs on the transcriptional level, we used the method of 4sU-tagging (17, 48-50). This method allows the metabolic labeling of newly transcribed RNA using 4-thiouridine (4sU), enabling the subsequent purification and separation of newly transcribed RNA from untagged pre-existing RNA (17, 49, 50). Differentiated macrophage-like THP-1 cells were treated with IFNα14 for 8 h or 24 h. 30 min before harvesting the cells, 4sU was added at a concentration of 500 µM. After purification and separation of the freshly transcribed and biotinylated RNA using Streptavidin-coated magnetic beads, changes in transcription rates were measured via RT-qPCR. The levels of newly transcribed *ISG15* mRNA were strongly enhanced after both 8 h and 24 h of treatment with IFNα14 (70-fold and 440-fold compared to untreated, respectively) (**Fig 4a**). In contrast, the levels of newly transcribed *SRSF1* mRNA were severely reduced after 8 h, with a reduction in relative mRNA expression levels of around 10-fold when compared to the control. After 24 h, *SRSF1* mRNA expression levels recovered but relative mRNA expression levels was still reduced by 2-fold when compared to the control (**Fig 4b**). This data indicates that IFN-mediated SRSF1 downregulation most likely occurs on the transcriptional level.

**Fig 4:**
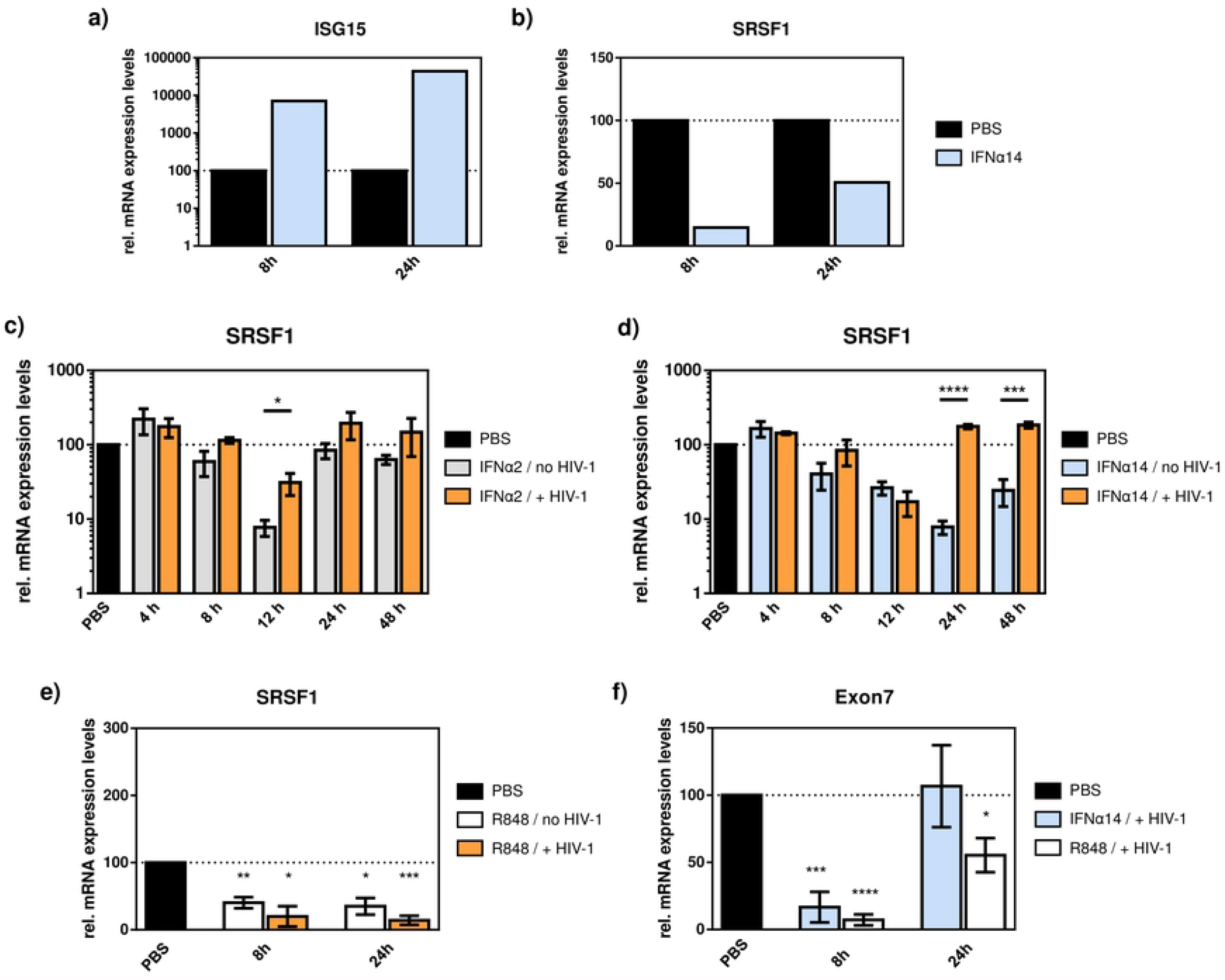
**a) – b) Changes in newly transcribed mRNAs upon treatment with IFNα14.** Differentiated THP-1 macrophages were treated with IFNα14 for 8 h or 24 h before labeling with 4sU for 30 min at a concentration of 500 µM. Cells were then harvested and total RNA isolated. Freshly transcribed RNA was labeled, purified and separated as described elsewhere (50). Relative mRNA expression levels of **a)** ISG15 and **b)** SRSF1 were measured via RT-qPCR. 2 biological replicates were pooled for qPCR analysis. GAPDH was used as loading control. **c) – d) HIV-1 counteracts repression of SRSF1 upon IFN-treatment**. Differentiated THP-1 macrophages were infected with the R5-tropic NL4-3 (AD8) at an MOI of 1. 16 h post infection, cells were treated with the indicated IFN subtype over a period of 48 h at a concentration of 10 ng/ml. Cells were then harvested, RNA isolated and subjected to RT-qPCR. Relative mRNA expression levels of SRSF1 in THP-1 cells after treatment with **c)** IFNα2 or **d)** IFN14. ACTB was used as loading control. Unpaired t-tests were calculated to determine whether the difference between the group of samples reached the level of statistical significance (* p<0.05, ** p<0.01 and *** p<0.001). **e) – f) TLR7/8 agonist R848 induces repression of SRSF1 mRNA expression levels**. Differentiated THP-1 macrophages were infected with the R5-tropic NL4-3 (AD8) at an MOI of 1 or mock infected. 16 h post infection, cells were treated with Resiquimod (R848) at a concentration of 30 µM for 8 h or 24 h respectively. Cells were then harvested, RNA isolated and subjected to RT-qPCR. **e)** Relative mRNA expression levels of SRSF1 after treatment with R848. **f)** Total viral mRNA levels were measured via RT-qPCR using a primer pair amplifying a sequence in Exon7. GAPDH was used as loading control. Unpaired t-tests were calculated to determine whether the difference between the group of samples reached the level of statistical significance (* p<0.05, ** p<0.01 and *** p<0.001).

### HIV-1 counteracts IFN-mediated repression of SRSF1

Since SRSF1 was identified as an IRepG in HIV-1 target cells (**Fig 3**), we were interested, whether an HIV-1 infection would influence the time-dependent downregulation. Therefore, we infected THP-1 macrophages with the R5-tropic HIV-1 laboratory strain NL4-3 (AD8) 16 h before IFN stimulation. Treatment with IFNα2, which led to a 13-fold downregulation in uninfected THP-1 cells after 12 h, only led to a 3-fold reduction of *SRSF1* mRNA upon HIV-1 infection (**Fig 4c**). In general, SRSF1 repression was more pronounced in IFNα2-treated uninfected cells compared to HIV-1 infected cells **(Fig 4c)**. IFNα14, which induced a long-lasting and 13-fold downregulation in non-infected cells, led to an overall weaker *SRSF1* mRNA repression of about 6-fold after 12 h in HIV-1 infected THP-1 cells (**Fig 4d**). Significantly higher SRSF1 expression levels were measured in IFNα14-treated HIV-1 infected cells after 24 h and 48 h when compared to non-infected cells (**Fig 4d**). These data suggest that HIV-1 counteracts the IFN-I-induced repression of SRSF1 in target cells, restoring somewhat balanced levels for efficient viral replication.

### HIV-1 sensing via TLR 7 and 8 is involved in the repression of SRSF1

TLR 7 and TLR 8 recognize single-stranded (ss) RNA and thus detect infections of RNA viruses such as HIV-1, leading to the secretion of cytokines like IFN-I (51, 52). Thus, we were further interested whether RNA sensing might play a role in the downregulation of SRSF1. Therefore, we tested the effect of the TLR 7/8 agonist Resiquimod (R848) on the expression level of *SRSF1* mRNA in HIV-1 infected or uninfected THP-1 macrophages.

Differentiated macrophage-like THP-1 cells were infected with the R5-tropic HIV-1 laboratory strain NL4-3 (AD8) or mock infected 16 h before treatment with R848 or IFNα14. The obtained results from the treatment of HIV-1 infected or mock infected THP-1 cells with IFNα14 were described in the previous section **(Fig 4d)**. Treatment with R848 led to a significant repression of *SRSF1* mRNA levels in uninfected cells after 8 h and 24 h (2.5- and 3-fold respectively) (**Fig 4e**). After 8 h, *SRSF1* mRNA levels were repressed by 5-fold while after 24 h even a 6-fold downregulation was detected in HIV-1 infected THP-1 cells (**Fig 4e**). Total viral mRNA levels were measured to investigate the impact of IFNα14 or R848 on viral replication. RT-qPCR was performed using a primer pair amplifying a sequence in Exon 7, which is present in all viral mRNA transcripts **(Fig 5)**. The amount of total viral RNA was reduced roughly by 6-fold after 8 h upon treatment with IFNα14, while after 24 h total viral mRNA levels were comparable to the untreated control (**Fig 4f**). Upon treatment with R848, a 14-fold reduction of total viral mRNA expression levels was detected after 8 h. After 24 h, the expression levels were still repressed by roughly 2-fold (**Fig 4f**). In conclusion, this data indicates that signaling pathways triggered by sensing via TLR 7 and 8 are involved in the repression of SRSF1, which might mediate the reduction of HIV-1 replication.

**Fig 5:**
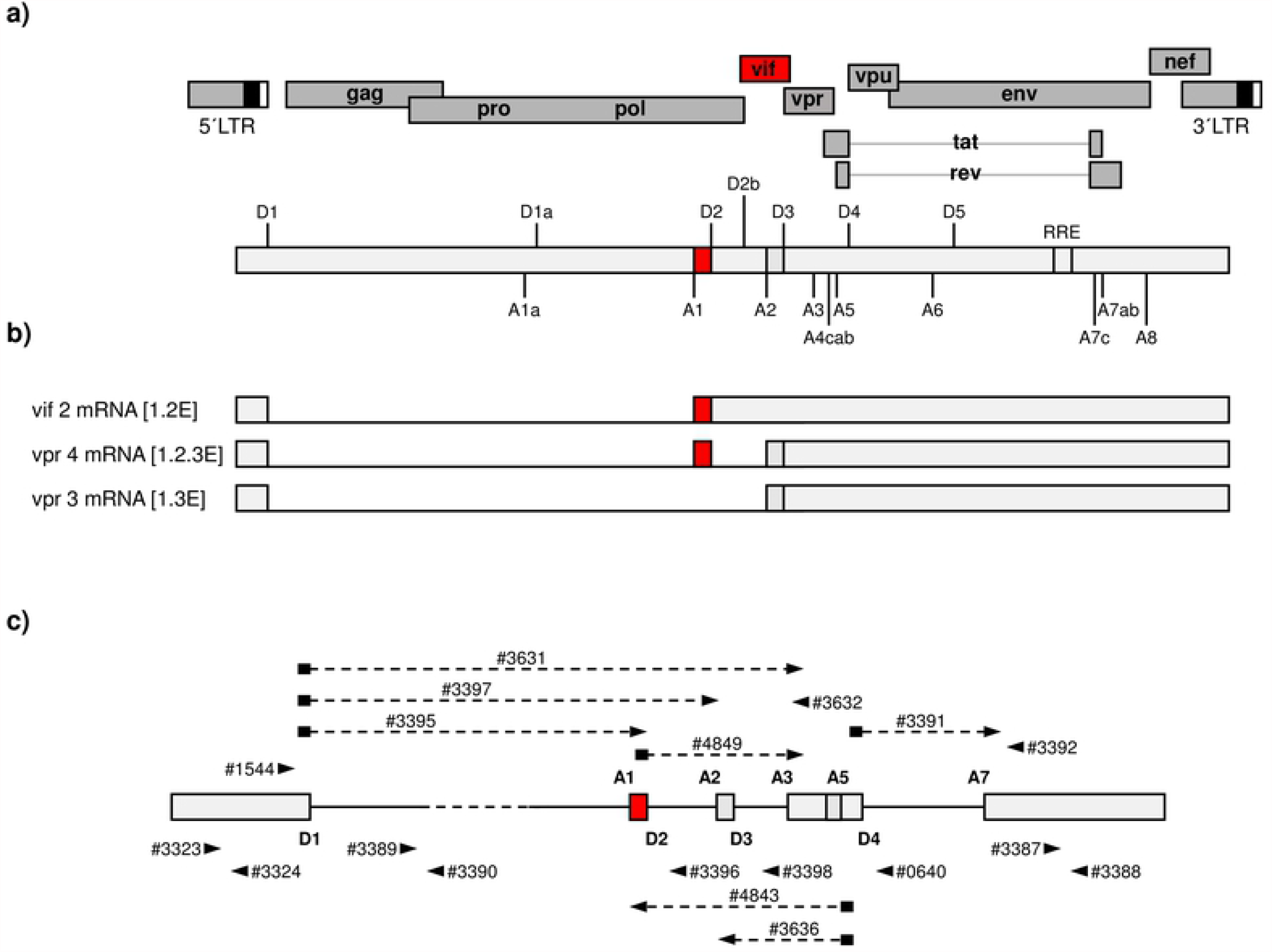
HIV-1 NL4-3 genome. **a)** HIV-1 genome with open reading frames (ORFs) and long terminal repeats (LTRs). 5’- and 3’-splice sites are indicated as well as the Rev response element (RRE). *Vif* Exon and ORF is highlighted in red. **b)** *Vif* and *Vpr* mRNs are spliced from 5’-ss D1 to 3’-ss A1 and 5’-ss D1 to 3’-ss A2 respectively, harboring the non-coding leader Exons 2 and 3. AUG-containing Introns 2 and 3 are contained respectively. **c)** Binding sites of primers for RT-qPCR and RT-PCR. Grey boxes indicate Exons, while straight lines indicate Introns. Black arrowheads indicate primers. Primers with black rectangle and black arrowhead connected via dashed line indicate Exon-junction primers.

### Knockdown of SRSF1 levels affect HIV-1 alternative splice site usage

Several binding sites of SRSF1 on the viral pre-mRNA have been identified (34, 36, 53), thus hinting at an important function of SRSF1 in HIV-1 RNA processing and replication. To evaluate the impact of IFN-mediated repression of SRSF1 on HIV-1 replication, we transiently silenced endogenous SRSF1 expression using a siRNA-based knockdown approach. After siRNA knockdown, HEK293T cells were transiently transfected with the HIV-1 laboratory strain pNL4-3 PI952 (54). 72 h post transfection, cells and virus-containing supernatant were harvested and subjected to various analyses.

Knockdown efficiency was verified via One-Step RT-qPCR, with siRNA inducing a knockdown of the SRSF1 gene expression of >80 % when compared to the negative control siRNA (**Fig 6a**). Intracellular total viral mRNA levels, measured via Exon 1 or 7 containing mRNAs which are present in all viral mRNA isoforms, were slightly elevated upon SRSF1 knockdown, albeit only significant for Exon 1 (**Fig 6b**).

**Fig 6:**
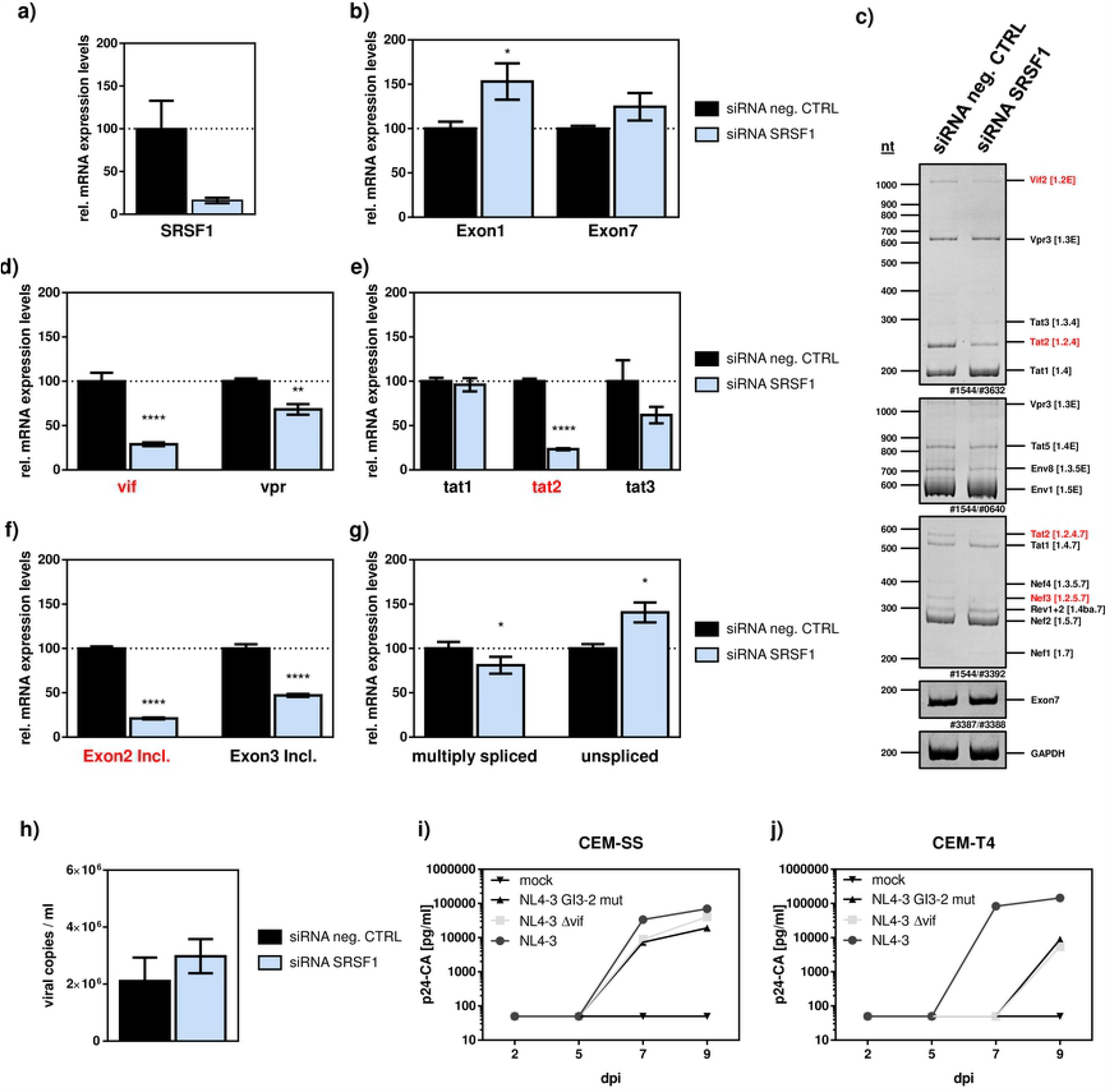
siRNA-induced knockdown of SRSF1. HEK293T cells were transfected with the proviral clone pNL4-3 PI952 (54) and the indicated siRNA. 72 h post transfection, cells were harvested and RNA and viral supernatant isolated. **a) – b)** Isolated RNA was subjected to RT-qPCR. Relative mRNA expression levels of **a)** SRSF1 and **b)** Exon 1 and Exon 7 containing mRNAs (total viral mRNA) normalized to GAPDH. **c)** Isolated RNA was subjected to RT-PCR using the indicated primer pairs for the 2 kb-, 4 kb- and *tat* mRNA-class. HIV-1 transcript isoforms are depicted on the right. To compare total RNA amounts, separate RT-PCRs amplifying HIV-1 exon 7 containing transcripts as well as cellular GAPDH were performed. PCR amplicons were separated on a 12% nondenaturing polyacrylamide gel and stained with Midori green Advance DNA stain (Nippon Genetics). **d) – g)** RT-qPCR results for relative mRNA expression levels of **d)** vif and vpr, **e)** tat1, tat2 and tat3, **f)** Exon 2 and Exon 3 containing and **g)** multiply spliced and unspliced mRNAs. HIV-1 mRNAs were analyzed using the indicated primers (**Table 1**). The splicing pattern of pNL4-3 PI952 was set to 100% and the relative splice site usage was normalized to total viral mRNA levels (Exon 7). Unpaired t tests were calculated to determine whether the difference between the group of samples reached the level of statistical significance (* p<0.05, ** p<0.01 and *** p<0.001). **h)** Cellular supernatant was used to determine viral copy number per ml. RT-qPCR was performed analyzing relative expression levels of exon 7 containing transcripts (total viral mRNA). **i)** CEM-SS and CEM-T4 cells were infected with wildtype NL4-3, NL4-3 Δ*vif*, NL4-3 G_I3_-2 mutant or mock infected. p24-CA ELISA of cellular supernatant was performed to determine virus production at the indicated time points.

Next, we analyzed the viral splicing pattern via semi-quantitative RT-PCR focusing on viral intron-less 2 kb-, intron-containing 4 kb- and *tat* specific mRNA-classes. SRSF1 knockdown resulted in significant alterations in the viral splicing pattern of all mRNA classes **(Fig 6c)**. These alterations could also be confirmed quantitatively by RT-qPCR using transcript specific primer pairs (**Fig 5, Fig 6d-g, Table 1**). The mRNAs of *vif* and *vpr*, the former of which is particularly crucial for efficient viral replication (55, 56), were significantly downregulated by 3- and 1.4-fold (**Fig 6d**). Since HIV-1 depends on the viral protein Vif to counteract APOBEC3G (A3G)-mediated antiviral activity of the host cell, this loss in *vif* mRNA might severely affect viral replication (14, 56, 57). While mRNA levels of *tat1* were not altered, both *tat2* and *tat3* mRNAs were repressed by 4- and 2-fold respectively (**Fig 6e**). Generally, the frequency of non-coding leader exons 2/3-including transcripts was significantly repressed by the factor of 5 and 2, respectively (**Fig 6f**). Since the levels of multiply spliced mRNAs (spliced from D4-A7) were slightly but significantly decreased by 1.25-fold, while levels of unspliced viral mRNAs (unspliced Intron 1) was significant increased by 1.4-fold (**Fig 6g**), knockdown of SRSF1 obviously shifts the ratio towards unspliced mRNAs.

**Table 1:**
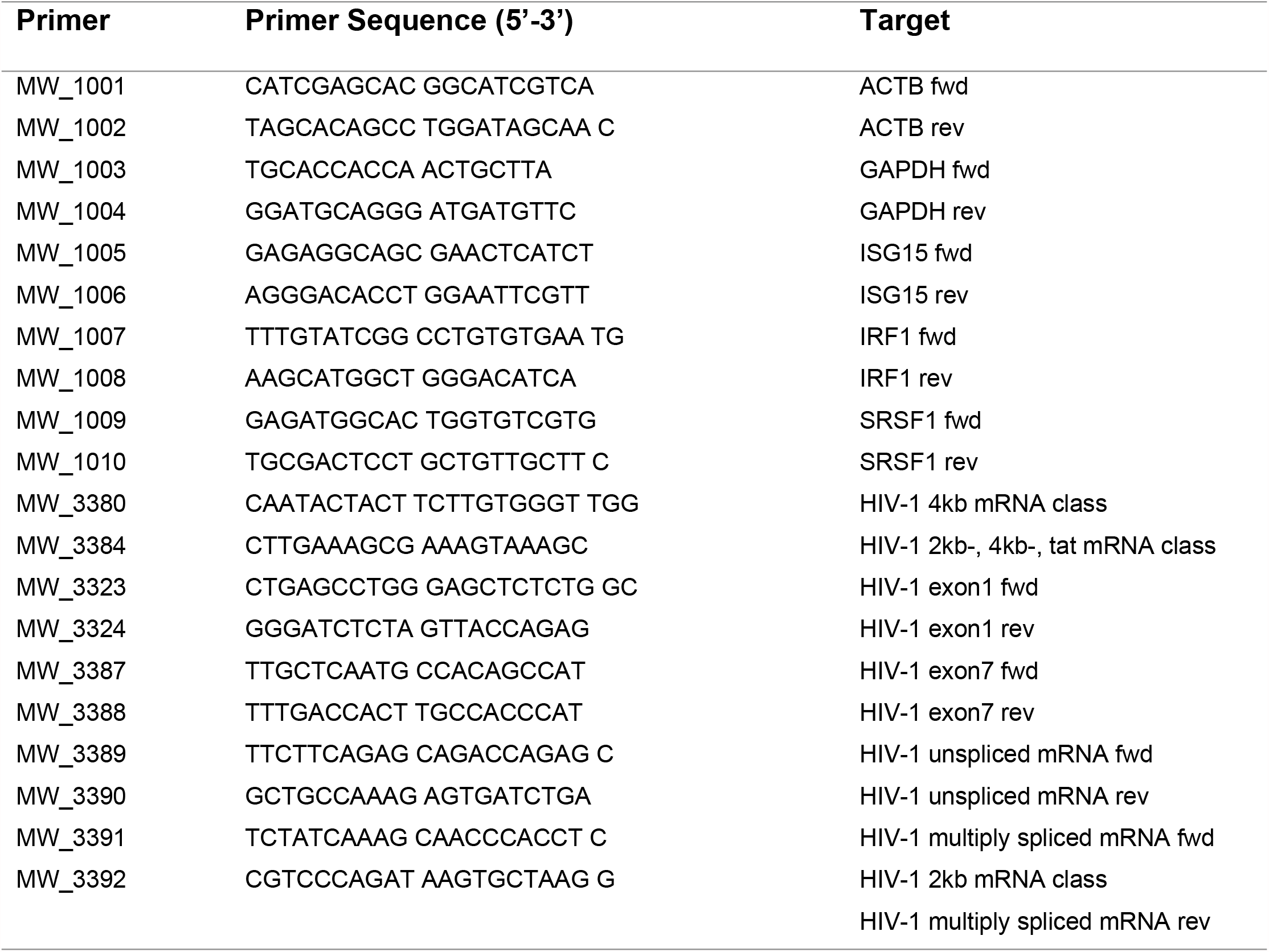
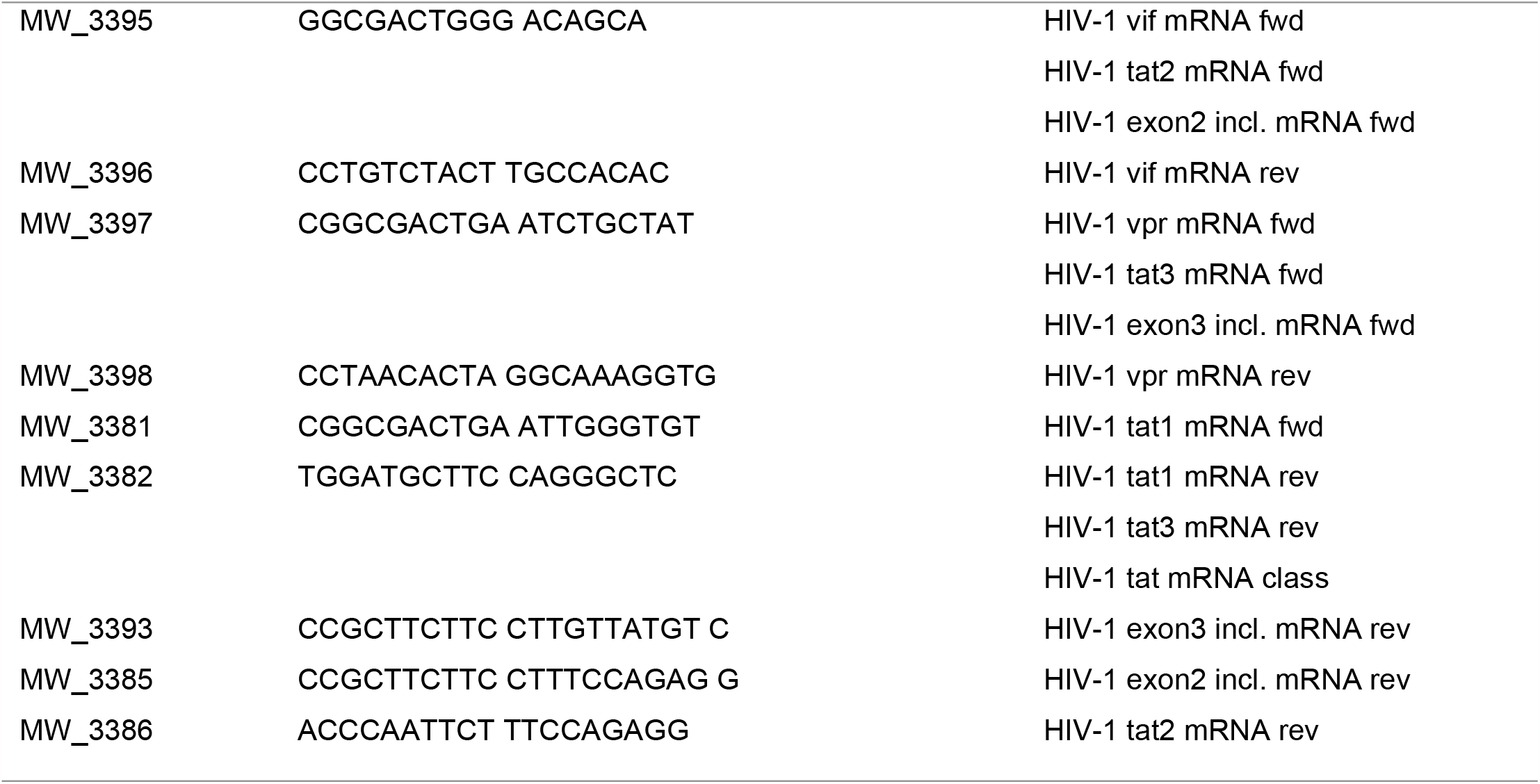
Primers used for RT-PCR and RT-qPCR.

Next, we were interested whether virus production would also be affected by the SRSF1 knockdown mediated changes in LTR transcription and alternative splicing. RT-qPCR analysis with viral RNA extracted from the cellular supernatant was performed, revealing a slight increase in viral copies, albeit not significant **(Fig 6h**). Since balanced levels of Vif are crucial for efficient viral replication in A3G-expressing cells (56, 58, 59), we were interested whether the significantly reduced levels of *vif* mRNA, caused by the siRNA-based knockdown of SRSF1, would impact viral replication capacity. Therefore, we performed replication kinetics in A3G-deficient CEM-SS cells (60, 61) and A3G-expressing CEM-T4 cells (62, 63) and monitored virus production by measuring p24 capsid protein production (p24-CA) in the cellular supernatant. As controls, we included NL4-3 wildtype virus, a *vif*-deficient NL4-3 Δ*vif (64)* and NL4-3 G_I3_-2 mut, which is characterized by reduced *vif* expression (60%) due to an inactivating mutation in the guanosine run element (G run) G_I3_-2 (58). As expected, in A3G low expressing cells (CEM-SS), both NL4-3 Δ*vif* and NL4-3 G_I3_-2 mut were able to replicate efficiently (**Fig 6i**). In A3G-expressing CEM-T4 cells, however, replication of both NL4-3 Δ*vif* and NL4-3 G_I3_-2 mut was strongly delayed indicating a less efficient viral replication capacity (**Fig 6j**). This data was in agreement with previously published data (58) suggesting that reduced levels of *vif* mRNA, as triggered by low SRSF1 amounts, strongly impair HIV-1 replication.

In conclusion, knockdown of SRSF1 disturbed the fine balance in the ratio of all HIV-1 mRNA classes and predominantly altered *vif* mRNA expression. Reduced Vif levels finally resulted in an impaired HIV-1 viral replication capacity in non-permissive cells.

### Overexpression of SRSF1 levels negatively affects HIV-1 replication

Furthermore, we were interested to which extend elevated SRSF1 levels would alter HIV-1 RNA processing. Therefore, we transiently transfected HEK293T cells with the HIV-1 laboratory strain pNL4-3 PI952 (54) and pcDNA-FLAG-SF2 (65). After 72 h, cells and virus containing supernatant were harvested and analyzed as described above.

Relative mRNA expression levels of *SRSF1* were enhanced by multiple orders of magnitude (**Fig 7a**) and protein expression and nuclear localization was confirmed using immune fluorescence microscopy (**S3 Fig**). As determined by the Exon 1 and 7 containing mRNAs, overexpressing SRSF1 resulted in a significant decrease in total viral mRNA levels (2-fold) (**Fig 7b**). To further analyze the effect of SRSF1 on the HIV-1 LTR promoter, cells were transiently transfected with a reporter plasmid harboring the firefly luciferase gene under the control of the HIV-1 LTR promoter (pTA-Luc-NL4-3). A plasmid coding for the viral transactivator Tat (pSVctat) (66) and a plasmid expressing SRSF1 (pEGFP-SF2) (67) were co-transfected. SRSF1 reduced the Tat-transactivated LTR promoter activity in a dose-dependent manner to 80% (0.1 µg) and 65% (0.2 µg) of the original activity respectively (**Fig 7c**).

**Fig 7:**
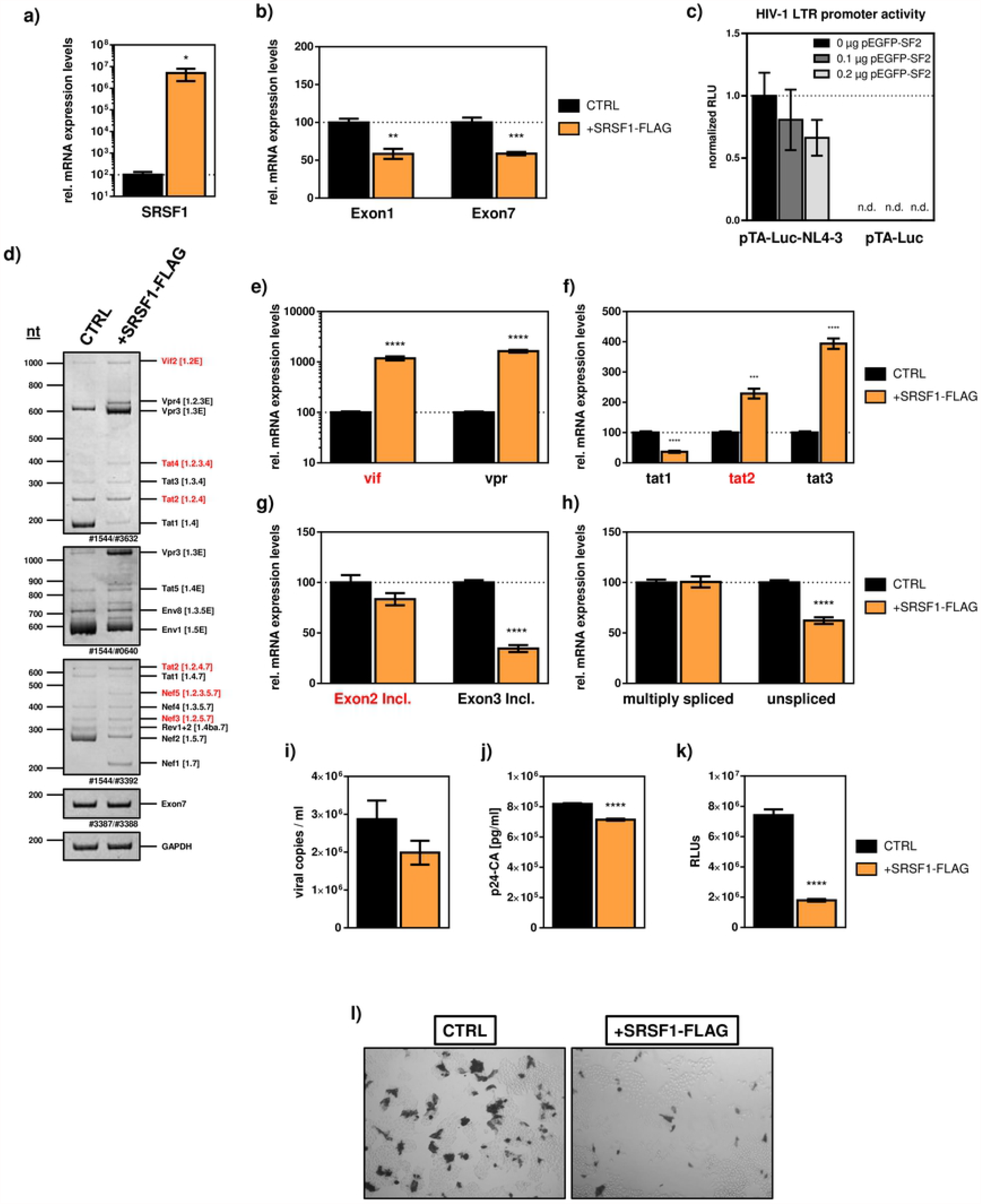
Overexpression of SRSF1. HEK293T cells were transfected with the proviral clone pNL4-3 PI952 (54) and pcDNA-FLAG-SF2 (65). 72 h post transfection, cells were harvested and RNA and viral supernatant isolated. **a) – b)** Isolated RNA was subjected to RT-qPCR. Relative mRNA expression levels of **a)** SRSF1 and **b)** Exon 1 and Exon 7 containing mRNAs (total viral mRNA) normalized to GAPDH. **c)** Vero cells were transiently co-transfected with pTA-Luc-NL4-3, pSVctat (66) and pEGFP-SF2 (67) at the indicated concentrations. Activity of HIV-1 LTR promoter was measured via luminescent read-out. **d)** Isolated RNA was subjected to RT-PCR using the indicated primer pairs for the 2 kb-, 4 kb- and *tat* mRNA-class. HIV-1 transcript isoforms are depicted on the right. To compare total RNA amounts, separate RT-PCRs amplifying HIV-1 exon 7 containing transcripts as well as cellular GAPDH were performed. PCR amplicons were separated on a 12% nondenaturing polyacrylamide gel and stained with Midori green Advance DNA stain (Nippon Genetics). **e) – h)** RT-qPCR results for relative mRNA expression levels of **e)** vif and vpr, **f)** tat1, tat2 and tat3, **g)** Exon 2 and Exon 3 containing and **h)** multiply spliced and unspliced mRNAs. HIV-1 mRNAs were analyzed using the indicated primers (**Table 1**). The splicing pattern of pNL4-3 PI952 was set to 100% and the relative splice site usage was normalized to total viral mRNA levels (Exon 7). Unpaired t tests were calculated to determine whether the difference between the group of samples reached the level of statistical significance (* p<0.05, ** p<0.01 and *** p<0.001). **i)** Cellular supernatant was used to determine viral copy number per ml. RT-qPCR was performed analyzing relative expression levels of exon 7 containing transcripts (total viral mRNA). **h)** Virus production was measured via p24-CA ELISA of cellular supernatant. **k) - l)** Viral infectivity was determined using TZM-bl reporter cells harboring the luciferase as well as the β-galactosidase expression cassette under the control of the HIV-1 LTR promoter. **j)** Measurement of luciferase activity. Unpaired t tests were calculated to determine whether the difference between the group of samples reached the level of statistical significance (* p<0.05, ** p<0.01 and *** p<0.001). **k)** X-Gal staining of TZM-bl cells incubated with cellular supernatant.

Viral splicing patterns were analyzed via semi-quantitative RT-PCR using primers specific for intron-less 2 kb-, intron-containing 4 kb- and *tat* specific mRNA-classes (**Fig 5, Table 1**). Overexpression of SRSF1 resulted in significant changes in the viral splicing pattern of all HIV-1 mRNA classes (**Fig 7d**). Alterations in the expression of HIV-1 specific mRNAs were also quantitatively confirmed by RT-qPCR using transcript specific primer pairs (**Fig 7e-h**). Levels of *vif* and *vpr* mRNA were increased by more than 10-fold (**Fig 7e**) while *tat1* mRNA expression was reduced by 3-fold and *tat2* and *tat3* mRNAs were upregulated by roughly 2- and 4-fold respectively (**Fig 7f**). In contrast to tat-specific mRNA-isoforms, the frequency of overall exon 2 inclusion was not altered upon SRSF1 overexpression. Exon 3 inclusion was reduced by roughly 3-fold (**Fig 7g**). The levels of multiply spliced mRNAs were not altered, while levels of unspliced viral mRNAs, measured via Intron 1-containing mRNAs, were significantly decreased by 1.6-fold (**Fig 7h**). Next, we performed RT-qPCR analysis with virus extracted from the cellular supernatant and found a decrease in viral copies, albeit not significant **(Fig 7i**). As determined by ELISA, the levels of p24 capsid were significantly lower when compared to mock transfected cells (**Fig 7j**). TZM-bl reporter cells were used to monitor infectivity of virus containing cellular supernatant harvested from transfected cells, revealing a significantly lower luciferase activity upon elevated SRSF1 levels (**Fig 7k**). This reduced infectivity was confirmed by X-Gal staining of TZM-bl cells infected with virus containing supernatants (**Fig 7l**).

Thus, overexpression of SRSF1 negatively affected Tat-LTR transcription and alternative splicing. Although *vif* mRNA levels, crucial for efficient viral replication, were significantly increased, both virus production and infectivity were significantly impaired.

## Discussion

Type I Interferons (IFNs) act as a first line of defense after viral infections (4, 5). Their mode of action includes the stimulation of ISGs including HIV-1 host restriction factors (11, 13), as well as the downregulation of IRepGs (16, 17), which are essential for viral replication. Together, both regulatory mechanisms establish an anti-viral state in the host cell. In this study, we were able to identify the cellular splicing factor SRSF1 as an IFN-I-repressed gene affecting HIV-1 post integration steps. For efficient viral replication, optimal SRSF1 levels are needed, which are in a narrow range.

SRSF1 was described as a key player in splicing regulation and gene expression of HIV-1 (33-35, 53, 68). Furthermore, SRSF1 was shown to have a much broader scope of action. Amongst a crucial role in cellular alternative splicing (26), SRSF1 regulates genome stability (69), translation (70), nuclear export (71) or the nonsense-mediated mRNA decay (NMD) pathway (72, 73). Loss of SRSF1 protein function resulted in G2 cell cycle arrest and induced apoptosis (74). Moreover, SRSF1 was defined as a proto-oncogene, since upregulation of SRSF1 favors the formation of a variety of cancers (75-77). Thus, the IFN-mediated downregulation of SRSF1 described in this manuscript might not only affect HIV-1 post integration steps, but also a variety of other cellular functions. We could show that the downregulation of SRSF1 upon IFN-treatment is time-dependent and after an initial repression, physiological levels are reached after different intervals depending on the cell type and IFN subtype. 4sU-labeling and isolation of newly transcribed RNA revealed the IFN-mediated repression of SRSF1 to result from an almost complete transcriptional shutdown of the SRSF1 gene. However, the exact mechanism how IFN-I induce this shutdown remains yet to be elucidated. A conceivable alternative or addition to a transcriptional shutdown could be the induction of an early RNA degradation mechanism. Since protein levels were reduced 12-24 h post RNA reduction, the role of host-mediated post-translational modifications (PTM) leading to protein degradation are unlikely. A prolonged SRSF1 downregulation is detrimental for a variety of cellular mechanisms and to guarantee balanced levels, SRSF1 was shown to maintain homeostasis through negative splicing feedback (29, 78). This autoregulatory mechanism will most likely also be responsible for the rapid upregulation that occurs immediately after the trough level of SRSF1 is reached.

Since expression levels of SRSF1 were repressed to a much higher extent in THP-1 macrophages than in Jurkat T-cells, the magnitude of SRSF1 repression seems to underlie cell type specific characteristics. Importantly, we were also able to confirm our cell culture derived results using primary cells.

During our initial screen, we investigated differences in the expression levels of SRSF transcripts between healthy and HIV-1-infected individuals. Upon HIV-1 infection, specific SRSF transcript levels and in particular SRSF1 were significantly lower in LPMCs and PBMCs when compared to healthy individuals (**Fig 1**). Since we have shown that IFN treatment has a direct effect on the downregulation of SRSF1, elevated IFN-I levels as a consequence of the HIV-1 induced chronic inflammation might play a key role.

Based on the increased ISG levels in acutely and chronically infected HIV-1 positive individuals, we confirmed both our patient cohorts as being representative. Furthermore, our findings suggest a direct correlation between the expression levels of ISGs including HIV-1 restriction factors and the expression levels of the cellular splicing factor SRSF1 in a physiological setting. In ART-treated patients, we observed that this difference could be reversed. A slight, non-significant increase was even observed, hence, currently we cannot exclude the possibility of ART-treatment having an influence on the transcript levels of SRSF1 or SRSF in general. Interestingly, it has been shown that more than 4000 genes are differentially expressed upon ART and that the IFN-induced JAK-STAT signaling pathway and several ISGs were downregulated following ART-treatment (79). However, whether this effect appears to be due to a decrease in inflammation or a direct effect of the administered substances needs further investigation.

While several IFNα subtypes elicit an antiviral activity suppressing HIV-1 infection, IFNα14 showed the most potent anti-HIV-1 activity of all subtypes both in PBMCs and LPMCs (7, 9). In an *in vivo* humanized mouse model, it was shown that combined treatment of ART and IFNα14 led to a more efficient suppression in HIV-1 plasma viral load (10, 80). While a clinical study to test a concomitant administration of ART and IFNα2 has been carried out (https://clinicaltrials.gov/ct2/show/results/NCT02227277), a benefit of subtype IFNα14 when compared to IFNα2 for a potential use in therapy could be a higher effectiveness and fewer side effects (81). The IFN-I-induced repression of SRSF1 expression was IFN subtype specific, with IFNα14 inducing the strongest downregulation of all IFNα subtypes. Furthermore, we suggest that SRSF1 repression is an IFN-I specific effect, since treatment with IFNγ only led to a negligible repression in THP-1 macrophages when compared to IFNα2 or IFNα14.

SRSF1 consists of two RRM, providing the RNA-binding specificity, and a relatively short RS-domain (25). The purine-rich pentamer GGAGA was identified as SRSF1 binding consensus motif via *in vivo* mapping (82, 83), with SRSF1 mainly binding in exonic splicing enhancer (ESE) regions although interestingly introns contain a high number of potential binding sites (25). Several binding sites of SRSF1 on the viral pre-mRNA have been identified (34, 36, 53) thus hinting at an important function of SRSF1 in HIV-1 RNA processing and replication.

Altered levels of SRSF1 have been shown to induce significant changes in HIV-1 LTR transcription (84). In agreement, we could confirm that overexpression of SRSF1 led to a significant reduction of total viral mRNA levels, while a siRNA-induced knockdown increased viral mRNA expression indicating a direct effect of SRSF1 on HIV-1 LTR transcription. Furthermore, alternative splicing was crucially affected by different SRSF1 levels. Both overexpression and knockdown led to significant alterations in the ratio of multiply spliced to unspliced mRNAs. Multiply spliced mRNAs are characterized by splicing from donor splice site D4 to acceptor splice site A7. The exonic splicing enhancer ESE3 is involved in the regulation of A7 usage and a known target of SRSF1 (35). Thus, altered levels of SRSF1 might affect splicing events occurring at A7. Inclusion of leader exons 2 and 3 significantly changed with altered levels of SRSF1, possibly indicating a direct effect of SRSF1 on splice acceptor A5. Both mRNA isoforms rely on the usage of A5, which is regulated by the bidirectional splicing enhancer and known target of SRSF1 ESE GAR (34).

While *vif* and *vpr* mRNA were significantly reduced upon SRSF1 knockdown, both mRNAs were strongly induced upon higher levels of SRSF1. Both accessory proteins Vif and Vpr play a crucial role in viral replication. While Vif counteracts the host restriction factor APOBEC3G enabling viral replication in non-permissive cells, Vpr is a multifunctional protein that amongst other purposes transports the pre-integration complex into the nucleus for viral integration or induces G2 cell cycle arrest, which enables the highest transcriptional activity of the HIV-1 LTR promoter (57, 85). *Vif* mRNA is spliced from D1 to A1, while *vpr* mRNA is spliced from D1 to A2 (3). Splice site A1 is regulated by the SRSF1-bound exonic splicing enhancer ESE M1/M2 (33). In presence of high SRSF1 levels, ESE M1/M2 facilitates the recognition of exon 2. Furthermore, cross-exon interactions between A1 and D2 (*vif* mRNA) and A2 and D3 (*vpr* mRNA) play a crucial role in the formation of the respective mRNAs (3, 58). In addition, further binding sites for SRSF1 on the HIV-1 pre-mRNA, as has been predicted *in silico* by the computational algorithm HEXplorer (86), might influence balanced HIV alternative splicing.

Interestingly, a HIV-1 infection counteracted the IFN-I-induced downregulation of SRSF1 for both treatments with IFNα2 and IFNα14. These findings hint to a crucial role of SRSF1 in HIV-1 replication and in particular post integration steps. Since treatment with TLR7/8 agonist R848 induced a similar mRNA expression pattern of SRSF1 than treatment with IFNα14, an involvement of viral sensing in the alteration of SRSF1 expression levels is indicated. TLR 7/8 signal through the MyD88-mediated IFN-regulatory factor (IRF) and NF-κB signaling pathways, stimulating the production of inflammatory cytokines and IFN-I (51, 87-91). Interestingly, the tendency of a concomitant HIV-1 infection to counteract SRSF1 repression upon treatment with IFNα14 could not be observed for the treatment with R848.

HIV infections through the mucosal route are frequently initiated by a single or a small quantity of transmitted founder viruses (TFV), which are relatively resistant to IFNs. Although IFN resistance has been linked to viral adaptations, specific viral properties that renders TFV IFN resistant is elusive. Roughly, 80% of all HIV-1 transmission events are established from a TFV (92). The genomic organization of HIV-1 TFV is generally comparable to the commonly used HIV-1 lab strain NL4-3 in terms of the used donor and acceptor splice sites. The usage of the specific splice sites however is strongly altered in many, but not all TFV (93). Here, analysis of the effect of altered levels of SRSF1 on LTR transcription and alternative splicing could give further insight into the ability of TFV to establish a successful HIV-1 transmission.

Reduced levels of *vif* mRNA, as observed upon knockdown of SRSF1, led to a significant impairment of virus replication. Elevated levels of SRSF1, which led to strongly increased levels of *vif* mRNA concomitantly reduced p24-CA levels. It has been also shown that high levels of Vif might inhibit viral infectivity by impaired proteolytic Gag precursor processing (94). In conclusion, balanced levels of Vif are required for efficient viral replication, which is in accordance to previously published data (55).

CEM-SS cells lack expression of A3G and several other host restriction factors, which counteract HIV-1 replication (56), and thus have a permissive phenotype allowing the replication of *vif*-deficient HIV-1 virus. The cell line CEM-T4 heterogeneously expresses A3G and acts as a semi-permissive cell line (62, 63) allowing both NL4-3 Δ*vif (64)* and NL4-3 G_I3_-2 mut to replicate, albeit with a strong delay in time when compared to the CEM-SS cells. Most likely, virus replication occurs in a subpopulation of cells that express no or low A3G levels.

In agreement with our data, SRSF1 in high concentrations was shown to block Tat-mediated LTR transcription by competing with the viral protein Tat for an overlapping binding sequence within the trans-activation response element (TAR) region. However, in the absence of Tat, SRSF1 increased the basal levels of HIV-1 transcription (40).

An interesting question that remains is whether drug targeting of SRSF1 would result in viral inhibition. The drug IDC16 was shown to block the replication of X4 and R5 tropic viruses, as well as clinical isolates via direct interaction with SRSF1 (95). The indole derivative can significantly influence splice enhancer activity of SRSF1 and impair splicing of HIV-1 pre-mRNA, thereby preventing the formation of multiple spliced mRNA isoforms and the expression of the early proteins Tat, Rev and Nef. However, because of the numerous influences on essential cellular processes, it is unlikely that such a drug will be used to treat HIV-1 infections.

In summary, our work shows that IFNs, in addition to the induction of antiviral genes, can also downregulate host factors which has a decisive influence on the early HIV-1 replication.

## Material and methods

### Cell culture, transient transfection and treatment

HEK293T, TZM-bl, Vero and ISRE-Luc reporter cells were maintained in Dulbecco’s modified Eagle medium (Gibco) supplemented with 10% (v/v) heat-inactivated fetal calf serum (FCS) and 1% (v/v) Penicillin-Streptomycin (P/S, 10.000 U/ml, Gibco). Jurkat, CEM-SS, CEM-T4 and THP-1 cells were maintained in Roswell Park Memorial Institute (RPMI) 1640 medium (Gibco) supplemented with 10% (Jurkat, CEM-SS and CEM-T4) or 20% (THP-1) (v/v) heat-inactivated fetal calf serum (FCS) and 1% (v/v) Penicillin-Streptomycin (10.000 U/ml, Gibco). THP-1 monocytes were treated with 100 nM 12-*O*-tetradecanoylphorbol-13-acetate (TPA) for 5 days to differentiate into macrophage-like cells. Differentiation was monitored via cell morphology and adhesion.

Transient transfection experiments were performed in six-well plates at 2.5 × 10^5^ HEK 293 T cells per well using TransIT^®^-LT1 transfection reagent (Mirus Bio LLC) according to the manufacturer’s instructions unless indicated.

IFNα subtypes were produced as previously described (7), IFNγ was purchased from PBL assay science (Piscataway, US). For the stimulation with IFN, 10 ng/ml of the respective IFN was added in fresh medium to the cells. The cells were then incubated at 37°C and 5% CO_2_ for the indicated amount of time before being harvested. Treatment with Resiquimod (R848) (Invivogen) was carried out at a final concentration of 30 µM for the indicated amount of time.

### RNA isolation, quantitative and semi-quantitative RT-PCR

The cells were harvested and total RNA was isolated using RNeasy Mini Kit (Qiagen) according to the manufacturer’s instructions. RNA concentration and quality was monitored via photometric measurement using NanoDrop2000c (Thermo Scientific). For reverse transcription (RT) 1 µg RNA was digested with 2 U of DNase I (NEB). After heat inactivation of the DNase at 70 °C for 5 min, cDNA synthesis for infection experiments was performed for 60 min at 50 °C and 15 min at 72 °C using 200 U SuperScript III Reverse Transcriptase (Invitrogen), 40 U RNase Inhibitor Human Placenta (NEB), 50 pmol Oligo d(T)23 (NEB) and 10 pmol Deoxynucleotide Triphosphate Mix (Promega). For all other experiments, cDNA synthesis was performed for 60 min at 42 °C and 5 min at 80 °C using ProtoScript II First Strand cDNA synthesis kit (NEB) according to the manufacturer’s instructions. Quantitative RT-PCR analysis was performed using Luna^®^ Universal qPCR Master Mix (NEB) and Rotor-Gene Q (Qiagen). Primers used for qPCR are listed in Table 1. ACTB or GAPDH were used as loading control for normalization. For qualitative analysis of HIV-1 mRNAs, PCR was performed using GoTaq G2 DNA Polymerase (Promega) according to the manufacturer’s instructions. PCR products were separated on non-denaturing polyacrylamide gels (12 %), stained with Midori green Advanced DNA stain (Nippon Genetics) and visualized with ADVANCED Fluorescence and ECL Imager (Intas Science Imaging).

### IFN-activity assay in RPE ISRE-luc reporter cell line

A reporter cell line of human retinal pigment epithelial (RPE) cells, stably transfected with a plasmid containing the firefly luciferase reporter gene under the control of the IFN-stimulated response element (ISRE), was used to determine the activity of the different IFNα-subtypes (7). Cells were seeded at 1.5 × 10^5^ cells per well in 12-well-plates and incubated overnight. The next day, cells were stimulated with 10 ng/ml of the respective IFNα subtype for 5 h. Cells were then lysed with Passive lysis buffer (Promega) and frozen at -80 °C overnight. After thawing, lysates were spun down and transferred to a white F96 Microwell plate (Nunc) before adding firefly luciferase substrate. Luminescent signal was measured using the GloMax^®^ Multi Detection System (Promega).

### Preparation of virus stocks, infection and replication kinetics

For the preparation of virus stocks, 6.5×10^6^ HEK293T cells were seeded in T175 flasks coated with 0.1% gelatin solution. The next day, cells were transiently transfected with 19 μg pNL4-3 or the respective proviral DNA using TransIT^®^-LT1 transfection reagent (Mirus Bio LLC) according to the manufacturer’s instructions. After 24 h the cells were supplemented with Iscove’s Modified Dulbecco’s Medium (IMDM, 10% (v/v) FCS, 1% (v/v) P/S) and incubated again overnight. The virus containing supernatant was then purified by filtration through 0.30 μm MACS SmartStrainers (Miltenyi Biotec), aliquoted and stored at -80 °C. Differentiated THP-1 cells and Jurkat cells were infected with the R5-tropic NL4-3 (AD8) (MOI, 1) or the dual tropic NL4-3 PI952 (MOI, 1) respectively with a spin-inoculation for 2 h at 1,500xg. 16 h post infection, indicated treatments were carried out. CEM-SS and CEM-T4 cells were infected as previously described (63). Virus production was monitored via p24-CA ELISA.

### Protein isolation and Western Blot

For protein isolation, cells were lysed with RIPA buffer (25 mM Tris HCl [pH 7.6], 150 mM NaCl, 1% NP-40, 1% sodium deoxycholate, 0.1% SDS, protease inhibitor cocktail [Roche]). The lysates were subjected to SDS-PAGE under denaturing conditions in 12% polyacrylamide gels using Bio-Rad Mini PROTEAN electrophoresis system (Bio-Rad). Gels were run for 90 min at 120 V in TGS-running buffer (25 mM Tris, 192 mM glycine, 0.1 % SDS (v/v)). NC-membrane (pore size 0.45 mm) was used for protein transfer using Bio-Rad Mini PROTEAN blotting system (Bio-Rad). Proteins were transferred for 1 h at 300 mA in transfer buffer (25 mM Tris, 192 mM glycine, 20% MeOH (v/v)). The membrane was blocked in TBS-T (20 mM Tris-HCl, 150 mM NaCl, 0.1 % Tween-20 (v/v) [pH 7.5]) with 5 % nonfat dry milk for 1 h at RT and then incubated overnight at 4°C with the primary antibody in TBS-T including 0.5 % nonfat dry milk. The membrane was washed three times for 10 min in TBS-T. The horseradish peroxidase (HRP) conjugated secondary antibody was added in TBS-T including 0.5 % nonfat dry milk and incubated for 1 h at RT. The membrane was washed 5 times for 12 min with TBS-T before ECL chemiluminescent detection reagent (Amersham) was added and read-out was performed with ADVANCED Fluorescence and ECL Imager (Intas). The following primary antibodies were used: Mouse antibody specific for SRSF1 (32-4500) from Invitrogen (Carlsbad, CA) and rabbit antibody specific for GAPDH (EPR16891) from Abcam (United Kingdom). The following horseradish peroxidase (HRP) conjugated secondary antibodies were used: anti-mouse HRP conjugate (315-035-048) from Jackson Immunoresearch Laboratories Inc. (West Grove, PA) and anti-rabbit HRP conjugate (ab97051) from Abcam (United Kingdom).

### p24-CA ELISA

For the quantification of HIV-1 p24-CA a twin-site sandwich ELISA was performed as previously described (55). Briefly, Immuno 96 MicroWell plates (Nunc) were coated with α-p24 polyclonal antibody (7.5 µg/ml of D7320, Aalto Bio Reagents) in bicarbonate coating buffer (NaHCO_3_, 100 mM, pH 8.5) overnight at room temperature. The plates were washed with TBS and blocked with 2 % non-fat dry milk powder in TBS for 1 h at room temperature. Empigen zwitterionic detergent (Sigma) was added to the samples for inactivation of HIV-1 and incubated for 30 min at 56 °C. Capturing of p24 and subsequent washing was carried out according to the manufacturer’s instructions (Aalto Bio Reagents). An alkaline phosphatase-conjugated α-p24 monoclonal antibody (BC1071 AP, Aalto Bio Reagents) was used for quantification of p24. Readout was performed with the Spark^®^ Microplate Reader (Tecan). Recombinant p24 was used to establish a p24 calibration curve.

### TZM-bl Luc assay and X-Gal staining

4,000 TZM-bl cells were seeded per well in 96-well plates and incubated overnight. 100 µl of supernatant was added to the cells and the plates were incubated for 48 h. For the luciferase assay, 50 µl lysis juice (p.j.k) was added after washing the plates with PBS and the plates were shaken for 15 min at room temperature. Next, the plates were frozen for at least 1.5 h at -80 °C before being thawed. Lysates were resuspended and transferred to a white F96 Microwell plate (Nunc) for luminescent readout. 100 µl beetle juice (p.j.k) was added per well and luminescence was measured with the Spark^®^ Microplate Reader (Tecan) at an integration time of 2 s. For the X-Gal staining, cells were washed with PBS and fixed in 0.06 % glutaraldehyde and 0.9 % formaldehyde for 10 min at 4 °C. Cells were washed twice with PBS and staining solution was added containing 400 mM K_3_[Fe(CN)_6_], 400 mM K_4_[Fe(CN)_6_], 100 mM MgCl_2_ and 20 mg/ml X-Gal. Cells were incubated overnight at 37 °C and overlayed with 50 % glycerol. Read-out was performed optically with light-microscopy.

### Measurement of HIV-1 replication kinetics

400,000 CEM-SS or CEM-T4 cells were infected with 1.6 ng of p24-CA of wildtype or mutant NL4-3 virus in serum-free RPMI medium (Invitrogen) at 37 °C. 6 h post infection, cells were washed and resuspended in complete RPMI medium. Aliquots of cell-free supernatant were harvested at indicated time points and p24-levels were measured via capture ELISA (see above).

### siRNA based knockdown

HEK293T cells were transiently transfected with the indicated siRNA at a final concentration of 8 nM using Lipofectamine 2000 (Thermo Scientific) according to the manufacturer’s instructions. The following siRNAs were used in this study: Silencer Select Negative Control #2 siRNA (Thermo Scientific) for the control siRNA and s12727 (Thermo Scientific) for SRSF1-specific siRNA.

### 4sU-tagging

Differentiated THP-1 cells were treated for 30 min with 4sU (Sigma Aldrich) at a final concentration of 500 µM for metabolic labeling of newly transcribed RNA following treatment with IFNα14 for the indicated amount of time. Labeling, purification and separation of freshly transcribed RNA was carried out as described elsewhere (50). Newly transcribed RNA concentration and quality was measured using NanoDrop2000c (Thermo Scientific).

### LTR-Luc plasmids

The LTR promoter of the HIV-1 laboratory strain pNL4-3 was cloned into the pTA-Luc backbone (Clontech) and is henceforth referred to as pTA-Luc-NL4-3. This plasmid encodes the firefly luciferase gene under the control of the cloned insert, allowing the measurement of the relative light units as direct correlation to the activity of the respective promotor. 100,000 Vero cells were seeded per well in 12-well plates and incubated overnight. Cells were then transiently transfected 1 µg pTA-Luc-NL4-3 and different amounts of pEGFP-SF2 using TransIT^®^-LT1 transfection reagent (Mirus Bio LLC) according to the manufacturer’s instructions. 24 h post transfection, cells were lysed using 350 µl Promega GloLysis buffer (Promega) One freeze and thaw cycle was performed before lysates were harvested using a rubber policeman and centrifuged at 13,000 rpm at 4 °C for 10 min. 50 µl of the cleared lysate was transferred to a white Nunc F96 Microwell plate (Nunc) for luminescent readout. 100 µl beetle juice (p.j.k.) was added per well luminescence was measured with the GloMax Discover (Promega) at an integration time of 10 s.

### PBMC isolation

Peripheral blood mononuclear cells (PBMCs) were isolated from whole blood samples by Ficoll density gradient centrifugation using LeucoSEP tubes (Greiner Bio-One) as described previously (96). RNA of isolated PBMCs was harvested as described above. This study has been approved by the Ethics Committee of the Medical Faculty of the University of Duisburg-Essen (14-6155-BO, 16-7016-BO, 19-8909-BO). Form of consent was not obtained since the data were analyzed anonymously.

### Statistical analysis

If not indicated differently, all experiments were repeated in three independent replicates. Statistical significance compared to untreated control was determined using unpaired student’s t-test. Asterisks indicated p-values as * (p<0.05), ** (p<0.01), *** (p<0.005) and **** (p<0.0001).

## Acknowledgements

We thank Christiane Pallas for excellent technical assistance. These studies were funded by the DFG (WI 5086/1-1; SU1030/1-2), the Jürgen-Manchot-Stiftung (H.S., M.W.), and the Medical Faculty of the University of Duisburg-Essen (H.S, K.S.). We thank Heiner Schaal for providing plasmid pSVctat (66) and Mirko Trilling for fruitful discussions. The following reagents were obtained through the AIDS Research and Reference Reagent Program, Division of AIDS, NIAID, NIH: TZM-bl cells from Dr. John C Kappes and Dr. Xiaoyun Wu. The following plasmids were obtained from Addgene: pEGF-SF2 (67) and pcDNA-FLAG-SF2 (65). The authors thank the Jürgen-Manchot-Stiftung for the doctoral fellowship of Helene Sertznig. This study has been approved by the Ethics Committee of the Medical Faculty of the University of Duisburg-Essen (14-6155-BO, 16-7016-BO, 19-8909-BO). Form of consent was not obtained since the data were analyzed anonymously. The funders had no role in study design, data collection and analysis, decision to publish, or preparation of the manuscript.

## References

1. Brass AL, Dykxhoorn DM, Benita Y, Yan N, Engelman A, Xavier RJ, et al. Identification of host proteins required for HIV infection through a functional genomic screen. Science. 2008;319(5865):921–6.

2. Stoltzfus CM. Chapter 1. Regulation of HIV-1 alternative RNA splicing and its role in virus replication. Advances in virus research. 2009;74:1–40.

3. Sertznig H, Hillebrand F, Erkelenz S, Schaal H, Widera M. Behind the scenes of HIV-1 replication: Alternative splicing as the dependency factor on the quiet. Virology. 2018;516:176–88.

4. Katze MG, He Y, Gale M, Jr. Viruses and interferon: a fight for supremacy. Nat Rev Immunol. 2002;2(9):675–87.

5. Lee AJ, Ashkar AA. The Dual Nature of Type I and Type II Interferons. Front Immunol. 2018;9:2061.

6. Gibbert K, Schlaak JF, Yang D, Dittmer U. IFN-alpha subtypes: distinct biological activities in antiviral therapy. Br J Pharmacol. 2013;168(5):1048–58.

7. Lavender KJ, Gibbert K, Peterson KE, Van Dis E, Francois S, Woods T, et al. Interferon Alpha Subtype-Specific Suppression of HIV-1 Infection In Vivo. J Virol. 2016;90(13):6001–13.

8. Sutter K, Dickow J, Dittmer U. Interferon alpha subtypes in HIV infection. Cytokine Growth Factor Rev. 2018;40:13–8.

9. Harper MS, Guo K, Gibbert K, Lee EJ, Dillon SM, Barrett BS, et al. Interferon-alpha Subtypes in an Ex Vivo Model of Acute HIV-1 Infection: Expression, Potency and Effector Mechanisms. PLoS pathogens. 2015;11(11):e1005254.

10. Sutter K, Lavender KJ, Messer RJ, Widera M, Williams K, Race B, et al. Concurrent administration of IFNalpha14 and cART in TKO-BLT mice enhances suppression of HIV-1 viremia but does not eliminate the latent reservoir. Scientific reports. 2019;9(1):18089.

11. Ivashkiv LB, Donlin LT. Regulation of type I interferon responses. Nature reviews Immunology. 2014;14(1):36–49.

12. Gao D, Wu J, Wu YT, D. F, Aroh C, Yan N, et al. Cyclic GMP-AMP synthase is an innate immune sensor of HIV and other retroviruses. Science. 2013;341(6148):903–6.

13. Stark GR, Kerr IM, Williams BR, Silverman RH, Schreiber RD. How cells respond to interferons. Annu Rev Biochem. 1998;67:227–64.

14. Doyle T, Goujon C, Malim MH. HIV-1 and interferons: who’s interfering with whom? Nature reviews Microbiology. 2015;13(7):403–13.

15. Okumura A, Lu G, Pitha-Rowe I, Pitha PM. Innate antiviral response targets HIV-1 release by the induction of ubiquitin-like protein ISG15. Proc Natl Acad Sci U S A. 2006;103(5):1440–5.

16. Megger DA, Philipp J, Le-Trilling VTK, Sitek B, Trilling M. Deciphering of the Human Interferon-Regulated Proteome by Mass Spectrometry-Based Quantitative Analysis Reveals Extent and Dynamics of Protein Induction and Repression. Frontiers in immunology. 2017;8:1139.

17. Trilling M, Bellora N, Rutkowski AJ, de Graaf M, Dickinson P, Robertson K, et al. Deciphering the modulation of gene expression by type I and II interferons combining 4sU-tagging, translational arrest and in silico promoter analysis. Nucleic acids research. 2013;41(17):8107–25.

18. Ocwieja KE, Sherrill-Mix S, Mukherjee R, Custers-Allen R, David P, Brown M, et al. Dynamic regulation of HIV-1 mRNA populations analyzed by single-molecule enrichment and long-read sequencing. Nucleic acids research. 2012;40(20):10345–55.

19. Kammler S, Leurs C, Freund M, Krummheuer J, Seidel K, Tange TO, et al. The sequence complementarity between HIV-1 5’ splice site SD4 and U1 snRNA determines the steady-state level of an unstable env pre-mRNA. RNA. 2001;7(3):421–34.

20. Freund M, Asang C, Kammler S, Konermann C, Krummheuer J, Hipp M, et al. A novel approach to describe a U1 snRNA binding site. Nucleic acids research. 2003;31(23):6963–75.

21. Smith CW, Chu TT, Nadal-Ginard B. Scanning and competition between AGs are involved in 3’ splice site selection in mammalian introns. Molecular and cellular biology. 1993;13(8):4939–52.

22. Barash Y, Calarco JA, Gao W, Pan Q, Wang X, Shai O, et al. Deciphering the splicing code. Nature. 2010;465(7294):53–9.

23. Wang Z, Burge CB. Splicing regulation: from a parts list of regulatory elements to an integrated splicing code. RNA. 2008;14(5):802–13.

24. Manley JL, Tacke R. SR proteins and splicing control. Genes Dev. 1996;10(13):1569–79.

25. Shepard PJ, Hertel KJ. The SR protein family. Genome Biol. 2009;10(10):242.

26. Zhou Z, Fu XD. Regulation of splicing by SR proteins and SR protein-specific kinases. Chromosoma. 2013;122(3):191–207.

27. Erkelenz S, Mueller WF, Evans MS, Busch A, Schoneweis K, Hertel KJ, et al. Position-dependent splicing activation and repression by SR and hnRNP proteins rely on common mechanisms. RNA. 2013;19(1):96–102.

28. Fu XD, Ares M, Jr. Context-dependent control of alternative splicing by RNA-binding proteins. Nature reviews Genetics. 2014;15(10):689–701.

29. Goncalves V, Jordan P. Posttranscriptional Regulation of Splicing Factor SRSF1 and Its Role in Cancer Cell Biology. Biomed Res Int. 2015;2015:287048.

30. Manley JL, Krainer AR. A rational nomenclature for serine/arginine-rich protein splicing factors (SR proteins). Genes & development. 2010;24(11):1073–4.

31. Krainer AR, Mayeda A, Kozak D, Binns G. Functional expression of cloned human splicing factor SF2: homology to RNA-binding proteins, U1 70K, and Drosophila splicing regulators. Cell. 1991;66(2):383–94.

32. Ge H, Manley JL. A protein factor, ASF, controls cell-specific alternative splicing of SV40 early pre-mRNA in vitro. Cell. 1990;62(1):25–34.

33. Kammler S, Otte M, Hauber I, Kjems J, Hauber J, Schaal H. The strength of the HIV-1 3’ splice sites affects Rev function. Retrovirology. 2006;3:89.

34. Caputi M, Freund M, Kammler S, Asang C, Schaal H. A bidirectional SF2/ASF- and SRp40-dependent splicing enhancer regulates human immunodeficiency virus type 1 rev, env, vpu, and nef gene expression. Journal of virology. 2004;78(12):6517–26.

35. Staffa A, Cochrane A. Identification of positive and negative splicing regulatory elements within the terminal tat-rev exon of human immunodeficiency virus type 1. Molecular and cellular biology. 1995;15(8):4597–605.

36. Jablonski JA, Caputi M. Role of cellular RNA processing factors in human immunodeficiency virus type 1 mRNA metabolism, replication, and infectivity. Journal of virology. 2009;83(2):981–92.

37. Jacquenet S, Decimo D, Muriaux D, Darlix JL. Dual effect of the SR proteins ASF/SF2, SC35 and 9G8 on HIV-1 RNA splicing and virion production. Retrovirology. 2005;2:33.

38. Ropers D, Ayadi L, Gattoni R, Jacquenet S, Damier L, Branlant C, et al. Differential effects of the SR proteins 9G8, SC35, ASF/SF2, and SRp40 on the utilization of the A1 to A5 splicing sites of HIV-1 RNA. J Biol Chem. 2004;279(29):29963–73.

39. Dillon SM, Guo K, Austin GL, Gianella S, Engen PA, Mutlu EA, et al. A compartmentalized type I interferon response in the gut during chronic HIV-1 infection is associated with immunopathogenesis. AIDS. 2018;32(12):1599–611.

40. Paz S, Krainer AR, Caputi M. HIV-1 transcription is regulated by splicing factor SRSF1. Nucleic Acids Res. 2014;42(22):13812–23.

41. Li Y, Sun B, Esser S, Jessen H, Streeck H, Widera M, et al. Expression Pattern of Individual IFNA Subtypes in Chronic HIV Infection. Journal of interferon & cytokine research : the official journal of the International Society for Interferon and Cytokine Research. 2017;37(12):541–9.

42. Deeks SG, Tracy R, Douek DC. Systemic effects of inflammation on health during chronic HIV infection. Immunity. 2013;39(4):633–45.

43. Zimmermann A, Trilling M, Wagner M, Wilborn M, Bubic I, Jonjic S, et al. A cytomegaloviral protein reveals a dual role for STAT2 in IFN-{gamma} signaling and antiviral responses. J Exp Med. 2005;201(10):1543–53.

44. Antonelli G, Scagnolari C, Moschella F, Proietti E. Twenty-five years of type I interferon-based treatment: a critical analysis of its therapeutic use. Cytokine Growth Factor Rev. 2015;26(2):121–31.

45. Billiau A, Matthys P. Interferon-gamma: a historical perspective. Cytokine Growth Factor Rev. 2009;20(2):97–113.

46. Bhat MY, Solanki HS, Advani J, Khan AA, Keshava Prasad TS, Gowda H, et al. Comprehensive network map of interferon gamma signaling. J Cell Commun Signal. 2018;12(4):745–51.

47. Pine R, Decker T, Kessler DS, Levy DE, Darnell JE, Jr. Purification and cloning of interferon-stimulated gene factor 2 (ISGF2): ISGF2 (IRF-1) can bind to the promoters of both beta interferon- and interferon-stimulated genes but is not a primary transcriptional activator of either. Mol Cell Biol. 1990;10(6):2448–57.

48. Melvin WT, Milne HB, Slater AA, Allen HJ, Keir HM. Incorporation of 6-thioguanosine and 4-thiouridine into RNA. Application to isolation of newly synthesised RNA by affinity chromatography. Eur J Biochem. 1978;92(2):373–9.

49. Windhager L, Bonfert T, Burger K, Ruzsics Z, Krebs S, Kaufmann S, et al. Ultrashort and progressive 4sU-tagging reveals key characteristics of RNA processing at nucleotide resolution. Genome Res. 2012;22(10):2031–42.

50. Garibaldi A, Carranza F, Hertel KJ. Isolation of Newly Transcribed RNA Using the Metabolic Label 4-Thiouridine. Methods Mol Biol. 2017;1648:169–76.

51. Heil F, Hemmi H, Hochrein H, Ampenberger F, Kirschning C, Akira S, et al. Species-specific recognition of single-stranded RNA via toll-like receptor 7 and 8. Science. 2004;303(5663):1526–9.

52. Meas HZ, Haug M, Beckwith MS, Louet C, Ryan L, Hu Z, et al. Sensing of HIV-1 by TLR8 activates human T cells and reverses latency. Nat Commun. 2020;11(1):147.

53. Tange TO, Kjems J. SF2/ASF binds to a splicing enhancer in the third HIV-1 tat exon and stimulates U2AF binding independently of the RS domain. Journal of molecular biology. 2001;312(4):649–62.

54. Polzer S, van Yperen M, Kirst M, Schwalbe B, Schaal H, Schreiber M. Neutralization of X4- and R5-tropic HIV-1 NL4-3 variants by HOCl-modified serum albumins. BMC Res Notes. 2010;3:155.

55. Widera M, Hillebrand F, Erkelenz S, Vasudevan AA, Munk C, Schaal H. A functional conserved intronic G run in HIV-1 intron 3 is critical to counteract APOBEC3G-mediated host restriction. Retrovirology. 2014;11:72.

56. Sheehy AM, Gaddis NC, Choi JD, Malim MH. Isolation of a human gene that inhibits HIV-1 infection and is suppressed by the viral Vif protein. Nature. 2002;418(6898):646–50.

57. Stopak K, de Noronha C, Yonemoto W, Greene WC. HIV-1 Vif blocks the antiviral activity of APOBEC3G by impairing both its translation and intracellular stability. Mol Cell. 2003;12(3):591–601.

58. Widera M, Hillebrand F, Erkelenz S, Vasudevan A, Munk C, Schaal H. A functional conserved intronic G run in HIV-1 intron 3 is critical to counteract APOBEC3G-mediated host restriction. Retrovirology. 2014;11(1):72.

59. Sheehy AM, Gaddis NC, Malim MH. The antiretroviral enzyme APOBEC3G is degraded by the proteasome in response to HIV-1 Vif. Nature medicine. 2003;9(11):1404–7.

60. Nara PL, Hatch WC, Dunlop NM, Robey WG, Arthur LO, Gonda MA, et al. Simple, rapid, quantitative, syncytium-forming microassay for the detection of human immunodeficiency virus neutralizing antibody. AIDS research and human retroviruses. 1987;3(3):283–302.

61. Nara PL, Fischinger PJ. Quantitative infectivity assay for HIV-1 and-2. Nature. 1988;332(6163):469–70.

62. Hache G, Harris RS. CEM-T4 cells do not lack an APOBEC3G cofactor. PLoS pathogens. 2009;5(7):e1000528.

63. Widera M, Erkelenz S, Hillebrand F, Krikoni A, Widera D, Kaisers W, et al. An Intronic G Run within HIV-1 Intron 2 Is Critical for Splicing Regulation of vif mRNA. Journal of virology. 2013;87(5):2707–20.

64. Mariani R, Chen D, Schrofelbauer B, Navarro F, Konig R, Bollman B, et al. Species-specific exclusion of APOBEC3G from HIV-1 virions by Vif. Cell. 2003;114(1):21–31.

65. Huang YQ, Ling XH, Yuan RQ, Chen ZY, Yang SB, Huang HX, et al. miR30c suppresses prostate cancer survival by targeting the ASF/SF2 splicing factor oncoprotein. Mol Med Rep. 2017;16(3):2431–8.

66. Schaal H, Pfeiffer P, Klein M, Gehrmann P, Scheid A. Use of DNA end joining activity of a Xenopus laevis egg extract for construction of deletions and expression vectors for HIV-1 Tat and Rev proteins. Gene. 1993;124(2):275–80.

67. Phair RD, Misteli T. High mobility of proteins in the mammalian cell nucleus. Nature. 2000;404(6778):604–9.

68. Tange TO, Damgaard CK, Guth S, Valcarcel J, Kjems J. The hnRNP A1 protein regulates HIV-1 tat splicing via a novel intron silencer element. The EMBO journal. 2001;20(20):5748–58.

69. Li X, Manley JL. Inactivation of the SR protein splicing factor ASF/SF2 results in genomic instability. Cell. 2005;122(3):365–78.

70. Sanford JR, Gray NK, Beckmann K, Caceres JF. A novel role for shuttling SR proteins in mRNA translation. Genes Dev. 2004;18(7):755–68.

71. Huang Y, Gattoni R, Stevenin J, Steitz JA. SR splicing factors serve as adapter proteins for TAP-dependent mRNA export. Molecular cell. 2003;11(3):837–43.

72. Aznarez I, Nomakuchi TT, Tetenbaum-Novatt J, Rahman MA, Fregoso O, Rees H, et al. Mechanism of Nonsense-Mediated mRNA Decay Stimulation by Splicing Factor SRSF1. Cell reports. 2018;23(7):2186–98.

73. Zhang Z, Krainer AR. Involvement of SR proteins in mRNA surveillance. Mol Cell. 2004;16(4):597–607.

74. Li X, Wang J, Manley JL. Loss of splicing factor ASF/SF2 induces G2 cell cycle arrest and apoptosis, but inhibits internucleosomal DNA fragmentation. Genes Dev. 2005;19(22):2705–14.

75. Fregoso OI, Das S, Akerman M, Krainer AR. Splicing-factor oncoprotein SRSF1 stabilizes p53 via RPL5 and induces cellular senescence. Mol Cell. 2013;50(1):56–66.

76. Das S, Anczukow O, Akerman M, Krainer AR. Oncogenic splicing factor SRSF1 is a critical transcriptional target of MYC. Cell Rep. 2012;1(2):110–7.

77. Karni R, de Stanchina E, Lowe SW, Sinha R, Mu D, Krainer AR. The gene encoding the splicing factor SF2/ASF is a proto-oncogene. Nat Struct Mol Biol. 2007;14(3):185–93.

78. Ding F, Su C, Chow K-HK, Elowitz MB. Dynamics and functional roles of splicing factor autoregulation. bioRxiv. 2020:2020.07.22.216887.

79. Massanella M, Singhania A, Beliakova-Bethell N, Pier R, Lada SM, White CH, et al. Differential gene expression in HIV-infected individuals following ART. Antiviral research. 2013;100(2):420–8.

80. Lavender KJ, Pace C, Sutter K, Messer RJ, Pouncey DL, Cummins NW, et al. An advanced BLT-humanized mouse model for extended HIV-1 cure studies. AIDS. 2018;32(1):1–10.

81. Sleijfer S, Bannink M, Van Gool AR, Kruit WH, Stoter G. Side effects of interferon-alpha therapy. Pharm World Sci. 2005;27(6):423–31.

82. Das S, Krainer AR. Emerging functions of SRSF1, splicing factor and oncoprotein, in RNA metabolism and cancer. Molecular cancer research : MCR. 2014;12(9):1195–204.

83. Chang B, Levin J, Thompson WA, Fairbrother WG. High-throughput binding analysis determines the binding specificity of ASF/SF2 on alternatively spliced human pre-mRNAs. Comb Chem High Throughput Screen. 2010;13(3):242–52.

84. Paz S, Caputi M. SRSF1 inhibition of HIV-1 gene expression. Oncotarget. 2015;6(23):19362–3.

85. Gonzalez ME. The HIV-1 Vpr Protein: A Multifaceted Target for Therapeutic Intervention. Int J Mol Sci. 2017;18(1).

86. Erkelenz S, Theiss S, Otte M, Widera M, Peter JO, Schaal H. Genomic HEXploring allows landscaping of novel potential splicing regulatory elements. Nucleic acids research. 2014.

87. Hemmi H, Kaisho T, Takeuchi O, Sato S, Sanjo H, Hoshino K, et al. Small anti-viral compounds activate immune cells via the TLR7 MyD88-dependent signaling pathway. Nat Immunol. 2002;3(2):196–200.

88. Zhou H, Yu M, Fukuda K, Im J, Yao P, Cui W, et al. IRAK-M mediates Toll-like receptor/IL-1R-induced NFkappaB activation and cytokine production. EMBO J. 2013;32(4):583–96.

89. Diebold SS, Kaisho T, Hemmi H, Akira S, Reis e Sousa C. Innate antiviral responses by means of TLR7-mediated recognition of single-stranded RNA. Science. 2004;303(5663):1529–31.

90. Ito T, Amakawa R, Kaisho T, Hemmi H, Tajima K, Uehira K, et al. Interferon-alpha and interleukin-12 are induced differentially by Toll-like receptor 7 ligands in human blood dendritic cell subsets. J Exp Med. 2002;195(11):1507–12.

91. Bender AT, Tzvetkov E, Pereira A, Wu Y, Kasar S, Przetak MM, et al. TLR7 and TLR8 Differentially Activate the IRF and NF-kappaB Pathways in Specific Cell Types to Promote Inflammation. Immunohorizons. 2020;4(2):93–107.

92. Joseph SB, Swanstrom R, Kashuba AD, Cohen MS. Bottlenecks in HIV-1 transmission: insights from the study of founder viruses. Nature reviews Microbiology. 2015;13(7):414–25.

93. Emery A, Zhou S, Pollom E, Swanstrom R. Characterizing HIV-1 Splicing by Using Next-Generation Sequencing. Journal of virology. 2017;91(6).

94. Akari H, Fujita M, Kao S, Khan MA, Shehu-Xhilaga M, Adachi A, et al. High level expression of human immunodeficiency virus type-1 Vif inhibits viral infectivity by modulating proteolytic processing of the Gag precursor at the p2/nucleocapsid processing site. The Journal of biological chemistry. 2004;279(13):12355–62.

95. Bakkour N, Lin YL, Maire S, Ayadi L, Mahuteau-Betzer F, Nguyen CH, et al. Small-molecule inhibition of HIV pre-mRNA splicing as a novel antiretroviral therapy to overcome drug resistance. PLoS pathogens. 2007;3(10):1530–9.

96. Widera M, Dirks M, Bleekmann B, Jablonka R, Daumer M, Walter H, et al. HIV-1 persistent viremia is frequently followed by episodes of low-level viremia. Med Microbiol Immunol. 2017;206(3):203–15.

